# Spatial transcriptome-metabolome integration reveals region-specific glial lipid dysregulation linked to Alzheimer’s pathology

**DOI:** 10.64898/2026.01.23.701345

**Authors:** Linfeng Xu, Hyunjun Yang, Julio Leon, Xiangpeng Li, Abby Oehler, Cyrus Modavi, Adam R. Abate, Carlo Condello

## Abstract

Alzheimer’s disease (AD) is associated with disrupted lipid metabolism, but how these abnormalities are organized across brain regions remains unclear. We developed iMIST, a spatial multi-omic approach that integrates metabolomic imaging, histology, and spatial transcriptomics within the same tissue section, and applied it to a mouse model combining progressive amyloid deposition and metabolic vulnerability. In plaque-rich gray matter, microglial activation was associated with Pld4 expression and glycerophospholipid remodeling. In contrast, plaque-sparse white matter showed oligodendrocyte-associated loss of Ugt8a expression and altered galactosylceramide metabolism. Both abnormalities intensified with age, indicating distinct but convergent forms of glial lipid dysregulation across anatomical compartments. These findings show that AD-associated metabolic dysfunction extends beyond plaque microenvironments and demonstrate how spatially integrated transcriptomic, metabolic, and histological measurements can reveal region specific mechanisms of glial vulnerability in neurodegeneration.

## Introduction

Alzheimer’s disease (AD) is a progressive neurodegenerative disorder marked by the accumulation of amyloid plaques and tau tangles, chronic neuroinflammation, and degeneration of vulnerable neuronal and glial populations.^1–5^ Multiple biological processes are known to contribute to its pathogenesis. In early-onset familial AD, mutations in *APP* or *PSEN* genes drive the accumulation of amyloid-β (Aβ), and amyloid plaques are sufficient to trigger downstream neurodegeneration. In contrast, the much more common sporadic, late-onset AD develops in the absence of such mutations and is instead associated with age-dependent decline and a constellation of metabolic, vascular, and inflammatory stressors.^6, 7^ Genetic studies have highlighted several late-onset risk alleles (*APOE ε4*, *ABCA7*, *CLU*, *TREM2*) that regulate lipid transport, signaling, and membrane turnover, pointing to lipid metabolism and glial cell function as central to disease susceptibility.^8–12^ Epidemiologic factors such as diabetes, vascular compromise, and chronic systemic inflammation further increase AD risk, suggesting that diverse insults converge on shared pathophysiological pathways.

A growing body of evidence potentially unites these observations under a single mechanistic framework. The glial-homeostasis hypothesis posits that AD arises from a progressive failure of glial lipid and energy regulation—the capacity of astrocytes, microglia, and oligodendrocytes to maintain metabolic balance and support neurons in the aging brain.^13–15^ Glial cells are uniquely sensitive to chronic activation: sustained inflammatory or metabolic stress transforms their normally protective functions into maladaptive states that impair lipid clearance, protein turnover, and energy production.^16, 17^ Over time, this loss of glial maintenance capacity may drive self-sustaining inflammation, accumulation of cellular debris, and neuronal dysfunction.^16–18^ In this view, amyloid plaques in early-stage disease could act as potent local stressors that exacerbate glial imbalance, while in late-stage disease, similar dysfunction may arise from diffuse metabolic and vascular stress.^19, 20^ In both cases, we postulate that the decisive event is not the presence of plaques per se but the collapse of glial homeostasis.

Microglia, the innate immune effectors of the brain, play central roles in neurodegenerative diseases of aging. Depending on disease stage and local pathology, they can be either protective or detrimental.^21, 22^ For example, microglia cluster around Aβ plaques, where they help contain aggregation^23, 24^, yet these same plaque-associated cells display aberrant metabolic demand^19, 20^, lipid droplet accumulation^18, 25, 26^, and proinflammatory activation that signal functional stress^27–29^. Beyond immune defense, microglia also regulate lipid turnover and maintain myelin integrity through interactions with oligodendrocytes and their progenitors^30–32^. Because these diverse metabolic and functional states emerge in distinct anatomical niches—around plaques, in white matter tracts, or near degenerating axons—understanding their mechanisms requires spatially resolved measurements that preserve tissue context. Systematic *in situ* approaches are therefore needed to localize and characterize dysregulated lipid metabolism in disease-activated microglia that may be central to AD pathogenesis.

This framework yields several testable predictions. Because glial stress responses are spatially localized, amyloid plaques and other focal lesions should induce microglial activation and lipid remodeling in their immediate surroundings, forming gradients of metabolic imbalance that decay with distance.^21–24^ White-matter tracts, which contain few plaques but rely on continual myelin renewal, should instead show oligodendrocyte lipid-synthesis deficits arising independently of amyloid.^33, 34^ Finally, as aging compounds metabolic and inflammatory stress, both types of dysfunction should intensify and expand in space, producing progressively broader signatures of glial and lipid dysregulation. Testing these predictions requires a method that can directly link transcriptional, metabolic, and pathological features within a unified spatial framework. Specifically, such an approach must capture all molecular layers within the same tissue manifold so that gradients of glial dysfunction can be quantified relative to anatomical and pathological landmarks. Although existing technologies can image lipids, profile metabolites, or map gene expression individually, no single workflow has yet united these modalities in one tissue section to resolve how local pathology, metabolic stress, and glial transcriptional states interact in space. The analytical framework must also account for the inherent noise and region-specific biases within spatial omics datasets. Conventional differential analysis treats each brain region as an independent experiment, which risks calling false-positive overlaps when a feature is significant in one comparison but behaves inconsistently in adjacent regions. A robust spatial multi-omic pipeline requires not only technical integration of metabolite imaging, histology, and transcriptomics on a single section, but also statistical cross-validation strategies that enforce biological consistency across spatial comparisons.

To meet this need, we developed iMIST (*in situ* Metabolomics, Imaging, and Sequencing in Tandem), a spatial multi-omic platform that integrates MALDI-based lipid and metabolite imaging, histology, and spatial transcriptomics in a single cryosection. Applying iMIST to the *APP23×APOEε4* mouse model of AD pathology and increased genetic risk, we found that cortical and thalamic regions with higher plaque loads exhibit signatures of microglial activation— including *Pld4*—that co-localize with amyloid deposits and glycerophospholipid remodeling. These results suggest that local pathological stress may amplify glial metabolic demand. In contrast, plaque-sparse white matter shows *Ugt8a* down-regulation and accumulation of galactosylceramides, indicating oligodendrocyte lipid-synthesis stress that arises distal to plaques. Across 6-, 12-, 15- and 18-month-old mice, both glial activation and lipid dysregulation intensified and expanded, matching a predicted temporal progression of glial failure. Together, these spatially resolved observations provide direct molecular support for the glial-homeostasis framework. Because iMIST operates on standard cryosections, it is compatible with both archived and fresh-frozen human brain tissue, enabling extension to clinical studies. iMIST thus delivers the first single-section, multi-modal datasets linking lipid metabolic state, histopathology, and glial gene expression in AD, establishing a platform for dissecting how local and global metabolic failure together shape neurodegeneration.

## RESULTS

### iMIST workflow to capture spatially resolved metabolomics and RNA-seq from a single brain section

To test whether Alzheimer’s pathology follows the spatial patterns predicted by the glial- homeostasis framework, we developed iMIST, an approach that directly links local pathology to glial molecular states *in vivo*. We required a model system that faithfully reproduced key features of disease pathology: the localized inflammatory stress associated with amyloid deposition and the broader metabolic vulnerability that impairs glial lipid handling. We generated a humanized *APP23*×*APOEε4* mouse line by crossing the established APP23 amyloid model^35^ with knock-in mice expressing human *APOEε4* on a murine *Apoe*-null background. The APP23 strain develops slowly progressive amyloid-β (Aβ) plaque deposition, a defining feature of AD. Accumulation begins around nine months of age, with moderate deposition across the cortex, hippocampus, and thalamus by twelve months, and continued expansion through eighteen months. The plaque phenotype plateaus near twenty-four months (the typical end of a laboratory lifespan). Introducing the known risk factor *APOEε4* into this background further humanizes the system by combining progressive amyloid deposition with impaired lipid transport and clearance, two factors thought to converge in driving loss of glial homeostasis. *APOEε4* exacerbates dysfunctional states in disease-associated microglia (DAMs) and worsens myelin phenotypes^36, 37^, which we hypothesize to sensitize glia to the chronic stress caused by amyloid.^38–42^ Together, these features create a model suited to examine how plaques and other regional stressors disrupt glial lipid metabolism and generate spatially patterned glial dysfunction across disease stages. For all experiments, *APOEε4* littermates lacking the *APP23* transgene served as “wildtype” (WT) controls.

The iMIST workflow builds on established commercial platforms, making it readily adoptable by other laboratories. As outlined in **Fig. 1**, the process begins with standard cryosectioning of brain tissue, and all data (metabolomic, histological, and transcriptomic) are collected from a single standard 14-µm section. This design preserves spatial relationships between Aβ plaques (5–50 µm in diameter) and surrounding cell populations (<20 µm), enabling precise molecular and anatomical alignment. For spatial metabolomic analysis, cryo-sectioned tissue is mounted onto a 10x Visium Spatial Gene Expression slide following the manufacturer’s instructions.^43, 44^ A conductive alumina MALDI membrane^45^ is then applied over the section and lightly sprayed with methanol. The membrane acts as both an absorptive matrix for metabolite capture and a conductive surface for ionization, effectively transforming the Visium slide into a MALDI-compatible matrix. A brief dry-ice-driven freeze–thaw cycle creates a thin condensation film between the tissue and membrane, facilitating efficient metabolite transfer. The prepared sample is then analyzed by MALDI-MSI^46, 47^ to map lipid and metabolite distributions. We deliberately set the MALDI-MSI raster step to 100 µm so that each pixel aligns exactly with the 100-µm resolution on the 10x Visium array, allowing facile spatial resolution registration between metabolite and transcriptomic measurements. This was supported by a two-zone definition of plaque area based on pixel counting (**Supplementary Fig. S1**).

**Fig. 1.**
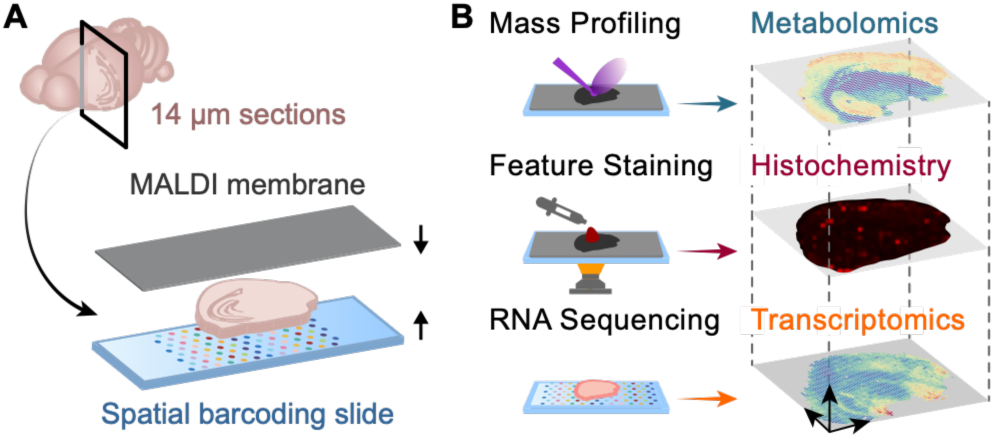
Schematic of multi-omics pipeline. **(A)** A cryo-sectioned brain slice is prepared and mounted on a 10x Visium gene expression slide before being layered with a conductive membrane. **(B)** The sample is then subjected to sequential measurements using MALDI-MSI, histology, and spatial transcriptomics

After MSI acquisition, the same section is rapidly stained with a red-shifted fluorescent amyloid-binding dye LDS722^48^ and imaged with an automated epi-fluorescence microscope to visualize Aβ plaques and cerebral amyloid angiopathy (CAA). The LDS722 dye shows strong signal over background (over 100-fold), enabling clear identification over background without further washing. Because the conductive membrane is opaque, imaging is performed through the transparent Visium glass slide, maintaining high spatial fidelity between plaque localization and brain architecture. Finally, spatial transcriptomic data are obtained from the same section using the standard Visium workflow. During this step, Proteinase K permeabilizes the membrane- sandwiched tissue to enable RNA capture and spatial barcoding. The earlier freeze–thaw step softens the membrane, allowing its clean removal without RNA loss. The resulting material is subjected to next-generation sequencing to generate spatially resolved gene-expression profiles.

The data analysis pipeline incorporates standard MSI and spatial transcriptomics analyses and requires no cross-sample registration or artifact correction because every dataset originates from a single cryosection. We developed a custom bioinformatic pipeline, D-iMISTer, to integrate and align pixels from each spatially-resolved dataset (fluorescent, MALDI IMS, and transcriptional data) to enable one to draw correlations and inferences from the combined data (**Supplementary Data S1**). Using D-iMISTer, we processed raw data from 18-month-old *APOEε4* control and *APP23×APOEε4* brain sections. The resulting multi-modal dataset (**Fig. 2A**) demonstrates precise alignment of transcriptomic, histological, and metabolomic information within the same tissue section (**Supplementary Data S2, S3**). The top panels of **Fig. 2A** display spatial transcriptomic gene distributions, the middle panels show fluorescently stained amyloid plaques, and the bottom panels present MALDI-MSI ion maps. The overlap among these layers confirms that iMIST achieves true spatial co-registration of histological, transcriptomic, and metabolomic information. Consistent with prior findings^49, 50^, ribonucleic acids remained largely intact through MALDI and histological processing, as assessed by spot-level unique molecular identifier (UMI) count concordance (**Supplementary Fig. S2, Supplementary Data S4**). The MALDI-MSI data showed that the conductive membrane effectively ionized molecules across the m/z 100–1000 range (**Supplementary Fig. S3**), making the method highly suitable for lipidomics studies using high-resolution MALDI instruments such as the Bruker Solarix 7.0T. Importantly, measurements of lateral diffusion by tracking two diagnostic lipid metabolites confirmed that there is minimal spatial misregistration of metabolites when using the membrane (**Supplementary Fig. S4**). Furthermore, advanced Alzheimer’s pathology was confirmed by the presence of LDS722-stained Aβ plaques and CAA in aged *APP23×APOEε4* brains. These are considered genuine based on morphological criteria and adjacent lipofuscins during manual segment plaque annotation. The results validate the technical robustness and spatial precision of the iMIST platform, confirming its ability to generate high-quality, co-registered metabolomic and transcriptomic data from a single brain section.

**Fig 2.**
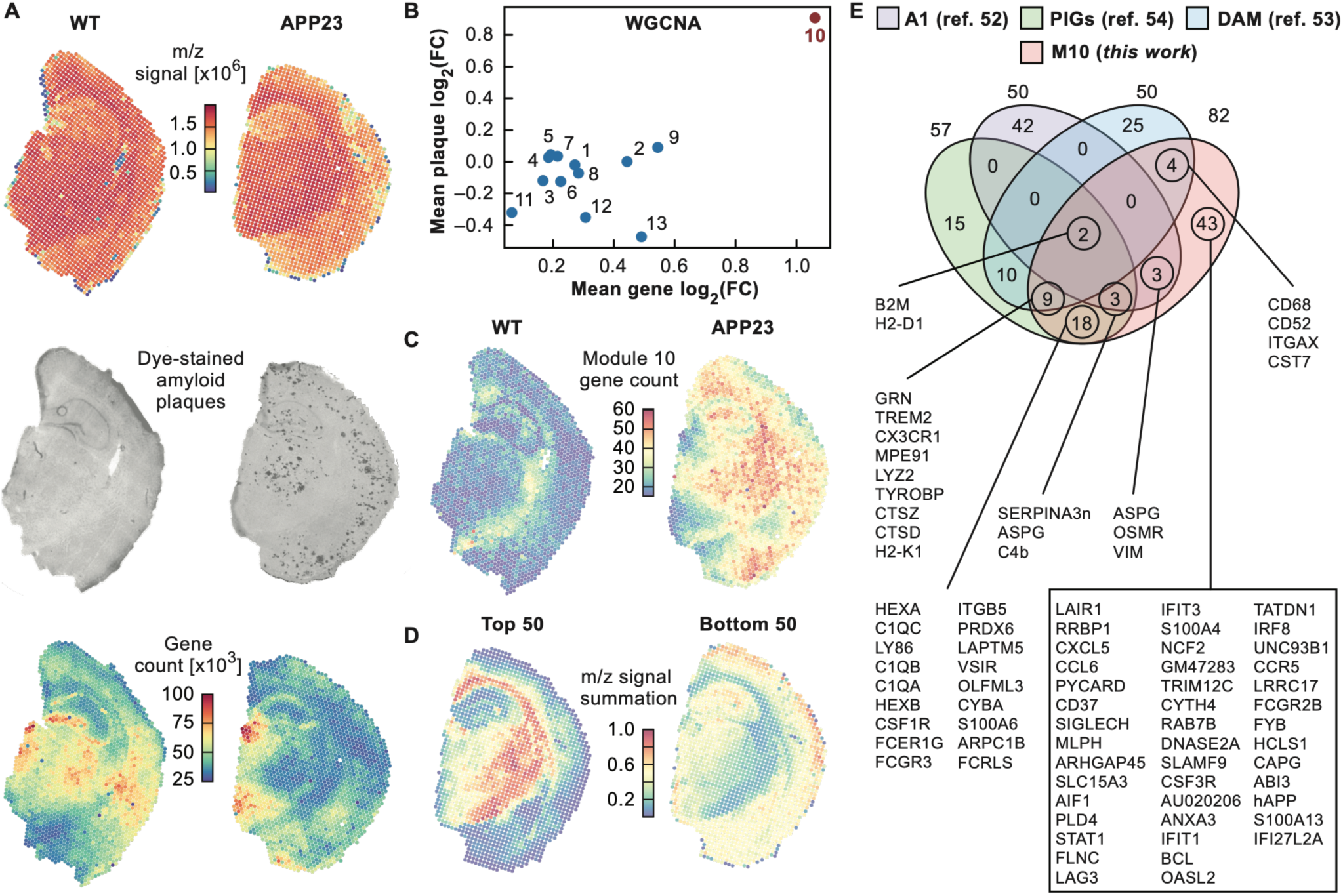
Illustrative multi-modal datasets obtained from iMIST. **(A)** Visualization of different iMIST datasets from whole brain slice of 18-months-old WT and APP23 mice. **(B)** Weighted Gene Co-expression Network Analysis (WGCNA) analysis reveals one module (designated “10”) as a significant cluster of genes highly correlated with plaque intensity. Mean plaque Log2(FC) represent the mean of the log-transformed Aβ index.^51^ **(C)** Module 10 gene expression signal within the whole brain slice of 18-months-old WT and APP23 mice. See **Supplementary Fig. S6** for the entire WGCNA and genes **(D)** Top 50 most correlated (left) and anti-correlated (right) m/z species within only 18-months-old APP23 brain slices. **(E)** Venn diagram illustrating how genes from module 10 (M10) overlap with previously reported AD-related gene studies. Beyond previously identified genes, our analysis identifies 43 novel gene candidates for future follow-up studies.

Beyond hardware integration, D-iMISTer introduces a key analytical innovation: rather than treating differential features from each brain region as independent discoveries, it can apply a consistency-supported cross-validation filter. A key metric is that features must meet consistent significance thresholds during primary comparison and simultaneous concordant fold-change direction in partner comparisons. This addresses the unique analytical challenges in spatial analysis: tissue heterogeneity, varying cell densities, and regional batch effects that can all produce significant but biologically inconsistent signals. The consistency-supported filter addresses this by leveraging the fundamental principle that a genuine biological response should propagate coherently across spatially connected regions, whereas technical artifacts are more likely to appear sporadically. By formalizing a reproducible statistical filter, D-iMISTer directly addresses a common pitfall in spatial omics where region-specific noise can generate apparently significant features that do not replicate across adjacent tissue areas. By enforcing cross-regional consistency, D-iMISTer distinguishes true spatial multi-omic signatures from artifacts, improving both the reliability of Venn overlap statistics and the biological interpretability of identified pathways. This framework is utilized across the datasets.

#### iMIST analysis captures region-specific and localized gene expression across the brain

It is well established that Alzheimer’s pathology is heterogeneous across the brain: gray matter regions such as cortex and thalamus develop dense amyloid plaques, whereas major white-matter tracts remain largely free of deposition. To confirm that our spatial data captured the expected brain region organization, we analyzed the spatial transcriptomic profiles to determine whether they could resolve major anatomical compartments and how these correspond to regional plaque distribution. Unsupervised Uniform Manifold Approximation and Projection (UMAP) of the spatial RNA-seq profiles identified distinct transcriptional programs corresponding to known brain regions (**Fig. 3A**), clearly separating cortex (Ctx), thalamus (TH), piriform cortical subplate (PIR_Ctx), lateral fiber tract (LFT), and corpus callosum (CC). These transcriptional clusters aligned with imaged anatomical and histological features, and matches the LDS722- identified concentration of plaques within the predominantly gray-matter regions (**Fig. 3B**), validating that the molecular data captured the expected anatomical and pathological organization of the brain, even after the multimodal acquisition workflow. Although the neuroanatomical atlas of different brain regions is well established, 10X Visium data provides a molecular segmentation of brain regions. This circumvents subtle differences that may be encountered when segmenting brain regions purely by histological methods (anatomical landmarks) and, thus, normalizing brain regions across different samples to improve the reproducibility of differential analysis.

**Figure 3.**
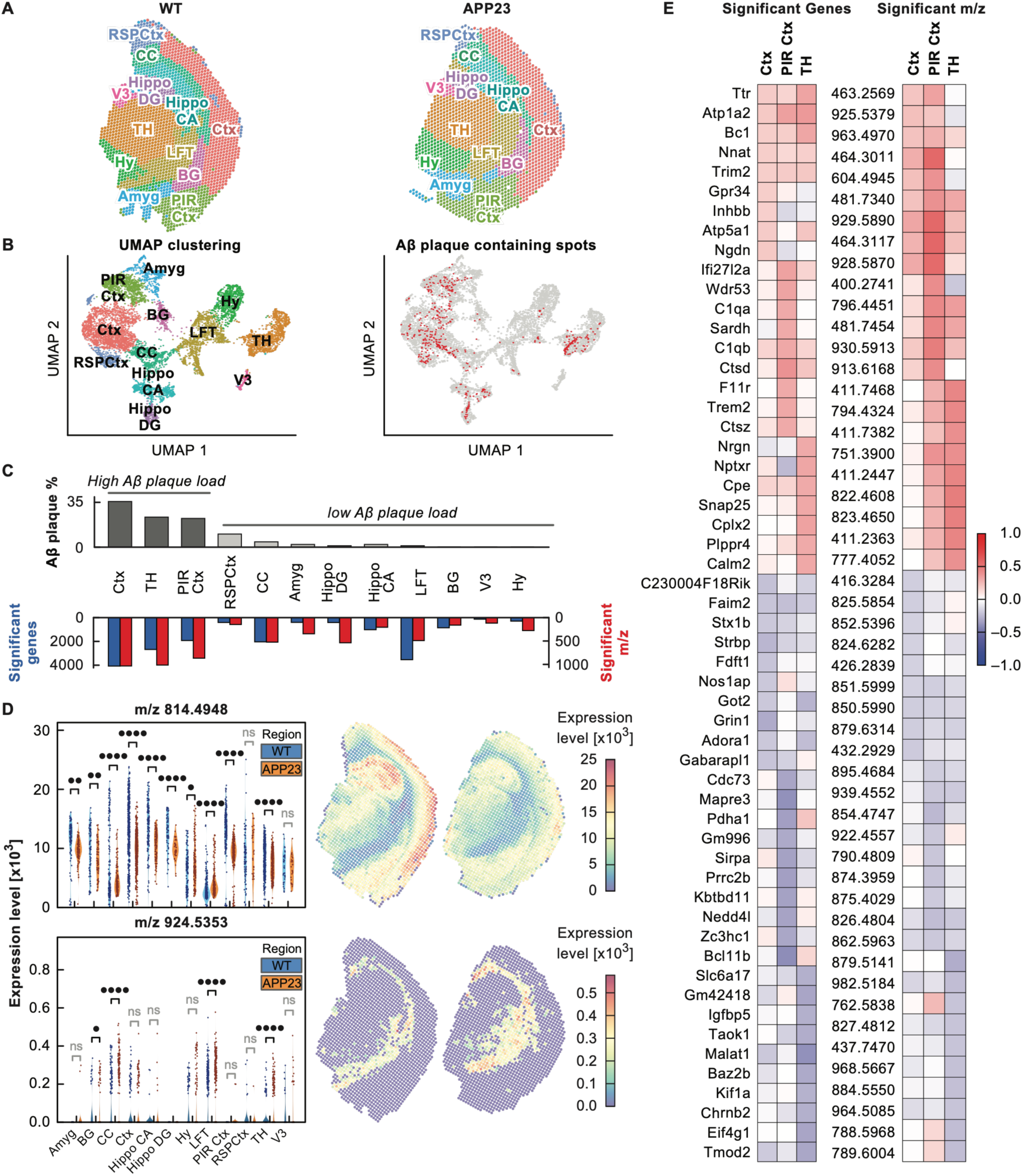
Multi-modal analysis obtained from D-iMISTer. **(A)** WT and APP23 brain section identified by unbiased UMAP clustering and classified based on histological location and gene expression signatures, mapped onto histology. Ctx, cortex; TH, thalamus; LFT, lateral fiber track; PIR Ctx, piriform cortical subplate; Hy, hypothalamus; CC, corpus callosum; Hippo_CA, hippocampus cornu ammonis; Amyg, amygdala; RSPCtx, retrosplenial cortex; Hippo_DG, hippocampus dentate gyrus; BG, basal ganglia; V3, ventricle 3. **(B)** UMAP plot of spatial transcriptomics and the plot of spatial distribution of Aβ plaques on the UMAP plot. **(C)** Histogram of DGE and DMA analysis representing the number of significant gene, m/z, as well as the number of spots containing Aβ plaques in a cluster. The entire DGE and DMA analysis is available in **Supplementary Data S2**. **(D)** Representative ion maps and violin plots for m/z 814.4948 and m/z 924.5353 illustrate their sharply contrasting spatial distributions and genotype-dependent abundance across WT and APP23 brains. Mann-Whitney U-Test was performed to compare the WT and APP23 ion counts. * p<=0.05, ** p <=0.01, *** p <=0.001, **** p <=0.0001. **(E)** Plaque centric integration of metabolomic and transcriptomic data ranks the 25 most strongly up- and down-regulated plaque-associated genes (PAGs) and metabolites (PAMs) within Aβ rich regions, revealing coordinated molecular alterations that whole-brain WGCNA could miss.

#### Plaques are associated with localized regions of transcriptomic and metabolic perturbations

Gray-matter regions are uniquely susceptible to amyloid deposition, which may reflect a property linked to their cellular composition and high metabolic demand. Plaques are not randomly distributed but instead form within these specific environments, suggesting that local factors—such as glial activation and lipid remodeling—both influence and are influenced by plaque formation. Microglia adopt specialized activation states near plaques, but whether these gene-expression changes coincide with local metabolic alterations has remained unclear. Because iMIST captures transcriptomic and metabolomic information from a single tissue section, it enables direct mapping of gene-expression modules to local lipid distributions with high spatial precision. Fluorescent plaque images from the same sections (**Fig. 2A**, middle row) provide the spatial reference for these analyses, allowing transcriptional and metabolic features to be correlated directly with histologically defined plaques.

We applied Weighted Gene Co-expression Network Analysis (WGCNA)^52^ to identify co-expressed gene networks distinguishing *WT×APOEε4* and *APP23×APOEε4* brains (**Fig. 2B**). Further, to ensure robust cross-comparison validation of differentially expressed features, we implemented a consistency-supported filtering strategy requiring each significant feature to pass the primary threshold in its own comparison (|avg_log2FC| > 0.25, adjusted p-value < 0.05) and exhibit concordant fold-change direction in the partner comparison (same-direction |avg_log2FC| > 0.25). This approach increased Jaccard overlap from 34.7% to 64.0% for up-regulated RNA features, from 20.4% to 60.5% for down-regulated RNA features, from 49.1% to 89.9% for up- regulated MS features, and from 50.3% to 83.5% for down-regulated MS features, substantially strengthening the reliability of identified spatial multi-omic signatures (**Supplementary Fig. S5; Supplementary Data S5**). Module 10 (82 genes) co-localized with Aβ plaques and was designated plaque-associated genes (PAGs). Consistent with this association, Module 10 strength increases with local plaque burden (**Fig. 2B–D; Supplementary Fig. S6**). The biological relevance of this module was supported by its overlap with established glial activation signatures, including A1^53^, disease-associated microglia (DAM)^22^, and previously reported plaque-induced gene (PIG) sets^51^ (**Fig. 2E**). Notably, this module of PAGs includes *Pld4* (phospholipase D family member 4), a lipid-metabolism enzyme expressed in disease-associated microglia and recurrently linked to plaque-proximal activation in Alzheimer’s models.^54, 55^

To determine whether transcriptional changes associated with plaques show corresponding metabolic alterations, we next analyzed the spatial distribution of metabolites. We first performed a pairwise correlation of Module 10 expression with all detected m/z signals. This analysis identifies metabolites whose abundance covaries with the spatial strength of the 82 PAGs program, revealing those that increase or decrease in concert with microglial activation. The resulting positive and negative associations defined a set of plaque-associated metabolites (PAMs) (**Fig. 2D** and **Supplementary Fig. S7**), identifying regions of high microglial activity that coincide with local remodeling of metabolite composition—predominantly lipid species, which dominate detection under our MALDI-MSI acquisition conditions.

To independently assess which metabolites are significantly altered between pathological and relatively unaffected regions, we next carried out differential metabolite-abundance analysis (p < 0.05, |log₂FC| > 50) on total-ion-normalized MALDI-MSI data. This comparison identified ions that varied significantly across regions, uncovering distinct metabolic signatures associated with plaque density. Integrating these metabolite changes with regional gene-expression data as a function of plaque burden revealed that the strongest combined perturbations occurred in Ctx, TH, and PIR_Ctx (**Fig. 3C**), whereas LFT and CC displayed comparable perturbations despite the lack of plaques. Spatial ion maps illustrated these regional patterns; for instance, m/z 814.4948 was nearly absent from myelin-rich LFT, whereas m/z 924.5353 was enriched in LFT and scarce elsewhere (**Fig. 3D**). Ranking the 25 most up- and down-regulated genes and metabolites within plaque-dense regions (**Fig. 3E**, expanded on in **Supplementary Fig. S8**) revealed spatially distinct molecular signatures that varied by region, reflecting localized differences suggestive of glial lipid dysregulation.

To determine which biochemical pathways are most affected by this localized glial lipid dysregulation, we integrated transcriptomic and metabolomic data at the pathway level. For each UMAP-defined region, we generated paired differential gene expression and differential metabolite abundance volcano plots (**Fig. 4A** and **Supplementary Data S2**) to visualize molecular changes associated with plaque density. Significant m/z features were tentatively annotated using the Human Metabolome Database^56^ within a 10-ppm mass window, accounting for [M+H], [M+Na], [M+K], and [M+H–H2O] adducts (instrument accuracy ≈ 900 ppb; see **Supplementary Data S3** for the full list of annotated m/z for putative metabolites and their isomers). Combining fold-change data from both datasets in MetaboAnalyst 6.0^57^, indicated glycerophospholipid metabolism as the most significantly perturbed pathway across plaque-dense regions, with the cortex showing nine distinct matched metabolites (**Fig. 4B** and **Supplementary Data S6**).

**Figure 4.**
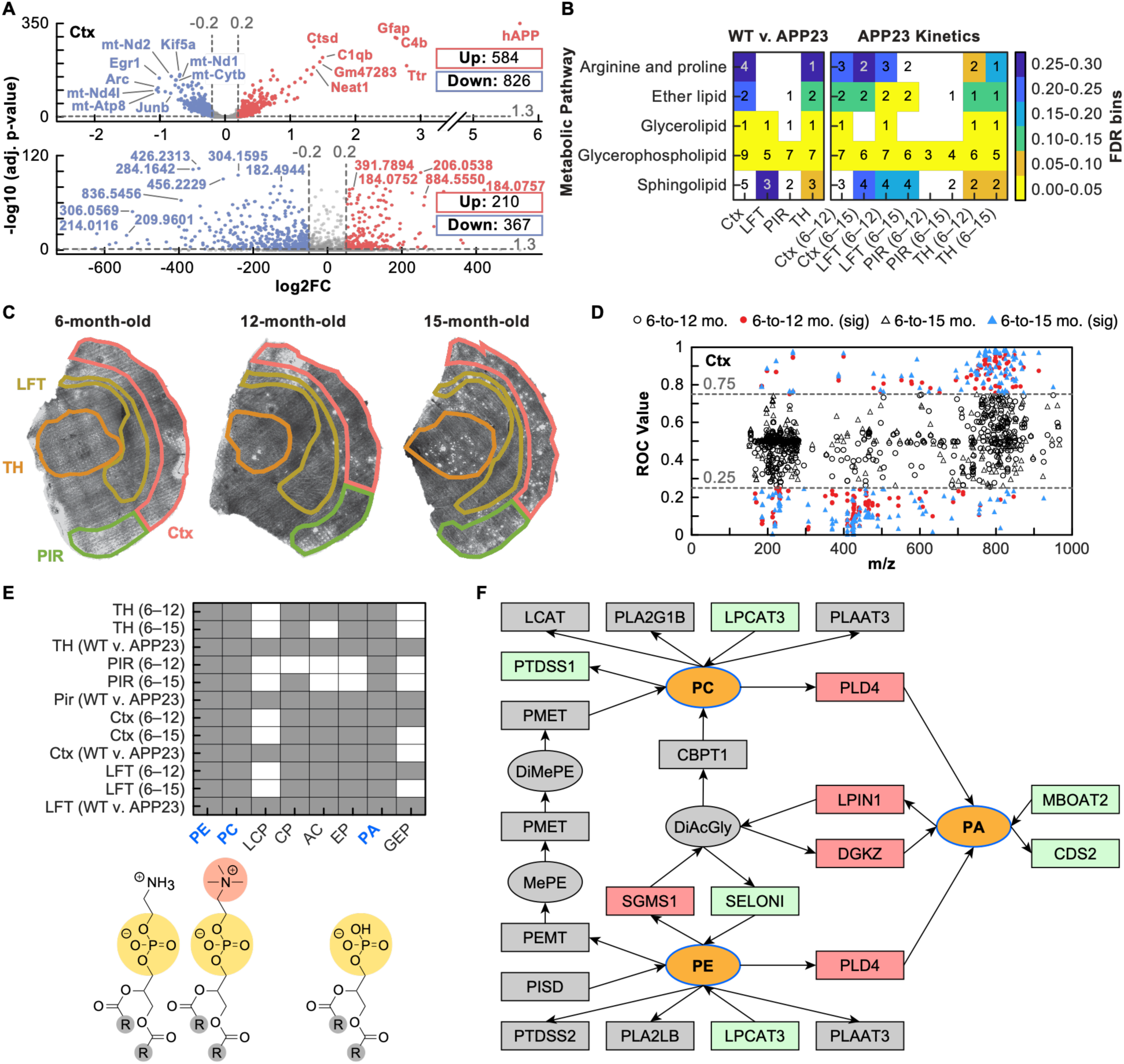
Integrated spatial metabolomic-transcriptomic analysis of APP23 brains. **(A)** Volcano plots from differential gene expression (DGE) and differential metabolite abundance (DMA) for the Ctx region. Gray dashed lines mark the fold change and False Discovery Rate (FDR) thresholds; insets enumerate examples of significant genes and metabolites. **(B)** KEGG pathway enrichment heatmaps produced with MetaboAnalyst. Left, WT vs APP23; right, APP23 kinetic data comparisons (6 months versus 12 or 15 months). Colors encode –log10 FDR, while the number within each cell provides a count of matched KEGG metabolites. **(C)** Epi-fluorescent images of APP23 coronal sections at 6, 12 and 15 months stained with LDS722 (Aβ plaques) and FluoroMyelin (white matter) after membrane-assisted MALDI imaging. Colored outlines indicate regions subjected to ROC analysis (Ctx, TH, PIR, LFT). **(D)** ROC plot for the Ctx. m/z features with AUC>0.75 (upregulated) or <0.25 (downregulated) are highlighted; red for 6-versus-12 months, blue for 6-versus-15 months; intermediate changes appear as hollow shapes. **(E)** Presence matrix of KEGG-annotated glycerophospholipid hits across region- and age-specific comparisons. Phosphatidic acid (PA), phosphatidylcholine (PC) and phosphatidylethanolamine (PE) are consistently altered. Corresponding structures depict the phosphate (yellow), headgroups (red), and alkyl tails (R group, gray). **(F)** Focused KEGG network of glycerophospholipid metabolism associated with PLD4 and relevant molecules. Ovals represent metabolites, rectangles genes; red = upregulated, green = downregulated in APP23 versus WT, gray = non-significant or undetected in iMIST.

Together, these results are in agreement with prior observations that gray matter regions undergoing glial homeostatic dysfunction exhibit localized amplification of plaque-induced metabolic stress, which are marked by microglial activation and glycerophospholipid remodeling.^22, 51, 58^ This combination likely reflects increased membrane-repair and lipid-clearance demand and defines a molecular environment that may promote plaque formation and sustain the feedback cycle of glial dysfunction. The spatial alignment of these signals with plaque location suggests the possibility of a distance-dependent relationship, in which metabolic stress and glial activation are strongest near plaques and quickly taper outward into surrounding tissue. Such spatial-dependence is consistent with recent findings of abnormal accumulation of lipid droplets in plaque-associated microglia but not in distal, non-plaque-associated microglia. ^25, 26^ Indeed, prior lipidomic and biochemical studies of AD models—performed largely on bulk cortical homogenates or low-resolution MS—have repeatedly implicated accelerated membrane glycerophospholipid turnover. These include shifts in phosphatidylcholine and phosphatidylethanolamine pools, enrichment of lysophospholipid metabolites, and dysregulated activities of phospholipase A2 and lysophospholipid acyltransferases that together drive lipid remodeling.^59–61^ These signatures have been interpreted as evidence for heightened membrane damage and compensatory repair—the type of processes that intersect with activated glia that upregulate lipid-handling programs in response to proteotoxic stress.^27, 28, 62^ However, because most prior lipid datasets average across heterogeneous cell populations and lack spatial context of plaques and transcriptomics and age-related lipid remodeling, they have not been able to determine whether these lipid changes are concentrated within plaque-rich niches versus broadly distributed across a region containing AD pathology. The results from iMIST affirms prior observations while providing deeper mechanistic insights to the cellular origins of biochemical alterations in selectively vulnerable brain regions.

#### Glycerophospholipid dysregulation intensifies with age and regional pathology

The glial-homeostasis framework predicts that aging progressively erodes the balance between lipid turnover, energy production, and inflammatory control, thereby amplifying lipid-metabolic stress in regions with the greatest pathological burden. To test whether these lipid pathways intensify with age, we profiled APP23 brains at 6, 12, and 15 months by first collecting MALDI MSI data. After MSI acquisition, we stained the same sections for myelin and amyloid plaques and imaged these markers to provide anatomical and pathological context, allowing all molecular readouts to be anchored to tissue landmarks (**Fig. 4C**). This workflow enabled us to evaluate how age-dependent lipid remodeling relates to both regional anatomy and evolving plaque pathology.

We quantified age-dependent shifts in the MSI data by performing Receiver Operating Characteristic (ROC) based contrasts (6→12 months and 6→15 months), flagging ions with Area Under the Curve (AUC) values > 0.75 or < 0.25 as significantly up- or down-regulated to identify m/z features that robustly increased or decreased over time. A representative cortical analysis illustrates the selection criteria and effect sizes (**Fig. 4D**). Across the age-related dataset, ions within the 400–1000 m/z range that corresponds primarily to lipid species exhibited substantially larger shifts than those in the small molecule metabolite window of 200–400 m/z window. By grouping 200–400, 400–600, and 600–1000 m/z intervals, the most pronounced remodeling with age was observed in the lipid-rich regions of the mass spectrum suggesting lipid metabolic reprogramming in APP23 mice with overt Aβ plaque load.

Significant ions were then queried against HMDB to generate tentative metabolite assignments, yielding a candidate list of metabolites for downstream interpretation. Mapping these putatively assigned metabolites suggested that glycerophospholipid metabolism might be a dominant age-associated signal in plaque-dense gray matter, with cortex showing multiple matched metabolites that increased with disease stage (**Fig. 4B, E**). Significant metabolites were analyzed jointly with significant genes from WT vs APP23 transcriptomic analysis. A Pld4-linked pathway was enriched that had association to the distributions of m/z species most likely corresponding to phosphatidic acid (PA, m/z 745.4610), phosphatidylcholine (PC, m/z 754.5194), and phosphatidylethanolamine (PE, 834.5624) (**Fig. 4F**). This provides a mechanistic rationale to focus subsequent analyses on tentatively assigned PA/PC/PE lipid remodeling across regions and age.

Spatial transcriptomics showed that Pld4, a prominent plaque-associated gene within this pathway, is progressively up-regulated in plaque-rich regions in the Ctx, PIR_Ctx, and TH (**Fig. 5A**). Higher resolution staining with RNAscope confirmed that Pld4 transcripts cluster around plaques and co-localize with microglial markers in a manner increasing from 6 to 15 months (**Fig. 5B**). Higher-magnification views found Pld4 transcript neighborhoods co-localizing with established microglial markers Csf1r and Clec7a specifically around plaque edges (**Fig. 5C**).

**Figure 5.**
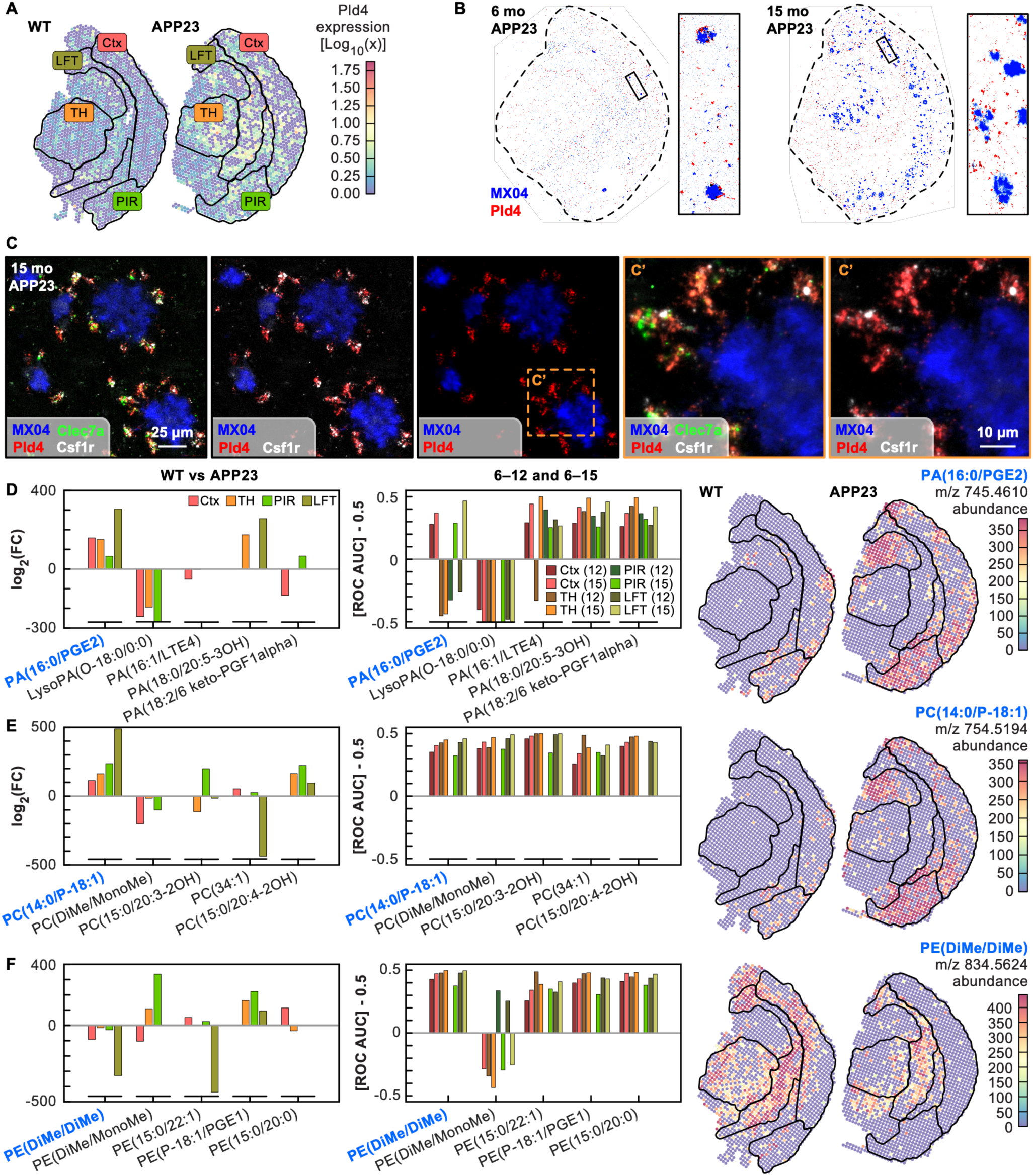
Spatiotemporal mapping of Pld4 expression and glycerophospholipid remodeling in APP23 brains. **(A)** Spatial transcriptomic heatmap of Pld4 (log_10_ normalized signal counts) showing widespread upregulation that co-localizes with Aβ plaque rich brain regions Ctx and TH. **(B)** RNAscope and tile-scan (20x) confocal microscopy analysis of 6-month- and 15- month-old APP23 brains reveals that Pld4 expression clusters around Aβ plaques. Pld4 is shown in red, while Aβ plaques are stained with the dye Methoxy-04 (MX04, blue) **(C)** At higher magnification (63x), Pld4 expression co-localizes with microglial markers Csf1r (gray) and Clec7a (green). **(D)** Phosphatidic acid (PA) isomers. Left, log2 fold change (APP23 vs WT) for the five most dysregulated PA species across cortex (Ctx, red), thalamus (TH, orange), piriform cortex (PIR, green) and lateral fiber tract (LFT, teal) at 6, 12 and 15 months. Center, corresponding ROC- AUC bars showing metabolite level changes with respect to AD pathology advancement. Right, ion count map for a representative PA species (m/z 745.4610) comparing WT and APP23 sections. **(E)** Phosphatidylcholine (PC) isomers analyzed using the same criteria, with spatial mapping of the significant ion m/z 754.5194. **(F)** Phosphatidylethanolamine (PE) isomers analyzed using the same criteria, with mapping of the significant ion m/z 834.5624. Histograms in D–F are ordered by the frequency with which each lipid was detected across all brain sections analyzed (WT vs APP23 and at 6, 12, and 15 months).

Tracking of isomer-resolved PA, PC, and PE species across regions reveals age-linked increases, decreases, and biphasic trajectories of these lipid families. **Figure 5D–F** presents spatiotemporal fingerprinting of glycerophospholipid remodeling in WT vs APP23 (left panels) versus APP23 brains by tracking isomer-resolved and significant PA (**5D**), PC (**5E**), and PE (**5F**) across four anatomically distinct regions (Ctx, TH, PIR, and LFT) at 6, 12, and 15 months. Here, we observed three recurrent types of age-dependent trajectories for metabolites in our time-course analysis: 1) monotonic increases with age, in which abundance rises progressively across the sampled intervals (consistent with gradual accumulation or escalating metabolic demand); 2) monotonic decreases with age, in which abundance steadily declines over time (consistent with depletion, reduced synthesis, or increased consumption); and 3) biphasic trajectories, in which the direction of change switches between early and late time windows (suggesting a transition in the underlying biology rather than a simple age effect).

Notably, these archetypes can be region-specific, such that the same metabolite can follow different trajectories depending on local pathology and cellular composition. For example, the ion species most corresponding to PA(16:0/PGE2) illustrates all three behaviors across anatomy: in Ctx, it shows a sustained progressive elevation from 6→12 and 6→15 months (a monotonic increase); in TH, it declines across 6→12→15 months within APP23 (a monotonic decrease, even though it can remain offset relative to WT at endpoint); and in PIR, it follows a biphasic pattern—downregulated early (6→12 months) when plaques are not yet evident, then upregulated later (6→15 months) coincident with emerging plaque pathology—consistent with a pathology-associated switch. Thus, by integrating both endpoint and time-course data, we identified distinct signatures of age-related metabolic remodeling, suggesting age-dependent plaque-associated lipid alterations.

#### White matter regions exhibit signatures of impaired oligodendrocyte lipid synthesis and aberrant galactosylceramide accumulation

Unlike gray matter, white matter rarely develops amyloid plaques despite being positioned as the functional bridge connecting grey matter loci, possibly because of its distinct cellular composition and metabolic environment. Nonetheless, extensive evidence shows that white matter deterioration—characterized by lipid-dependent changes such as myelin thinning, membrane fragmentation, and axonal rarefaction—occurs throughout the course of Alzheimer’s disease, beginning before plaque formation and worsening as pathology advances. White matter is dominated by glial cells, especially oligodendrocytes, which synthesize and maintain myelin, making them particularly sensitive to lipid metabolic stress. Consistent with this organization, we observe oligodendrocyte lipid-synthesis impairment (based on Ugt8a decrease) accompanied by selective accumulation of galactosylceramide (GalCer). reflecting manifestation of glial homeostatic dysfunction in this compartment. The GalCer lipid is a major myelin component that stabilizes the multilamellar sheath and supports continuous membrane renewal; its dysregulation is a hallmark of oligodendrocyte stress and early demyelination. Among regions largely devoid of Aβ plaques, the heavily myelinated white-matter bundle of the LFT, exhibited the strongest gene-expression and metabolic remodeling despite minimal amyloid pathology (**Fig. 3C**).

Pathway enrichment of LFT-specific m/z signals implicated GalCer metabolism, a network involving Ugt8a (GalCer synthase) and modifying enzymes Galc, Gal3st1, and Arsa (**Fig. 6A-D**). These enzymes together maintain sphingolipid balance in myelin membranes. Within this pathway, MALDI-MSI revealed a prominent ion at m/z 766.5554, assignable to the myelin lipid GalCer(d18:1/18:0), whose abundance increased sharply and specifically within the LFT (**Fig. 6E- G**). Longitudinal analysis further putatively identified an unsaturated analogue, GalCer(d18:1/18:1), following a similar white-matter-restricted trajectory, while most other ceramide species remained stable or declined over time (**Fig. 6H**), highlighting how iMIST can identify evidence of altered cellular metabolic states. The finding of a spatially confined lipid-remodeling signature in white matter centered on GalCer species and oligodendrocyte-associated expression changes, suggest that glial homeostatic stress plays a major role in this compartment.

**Figure 6.**
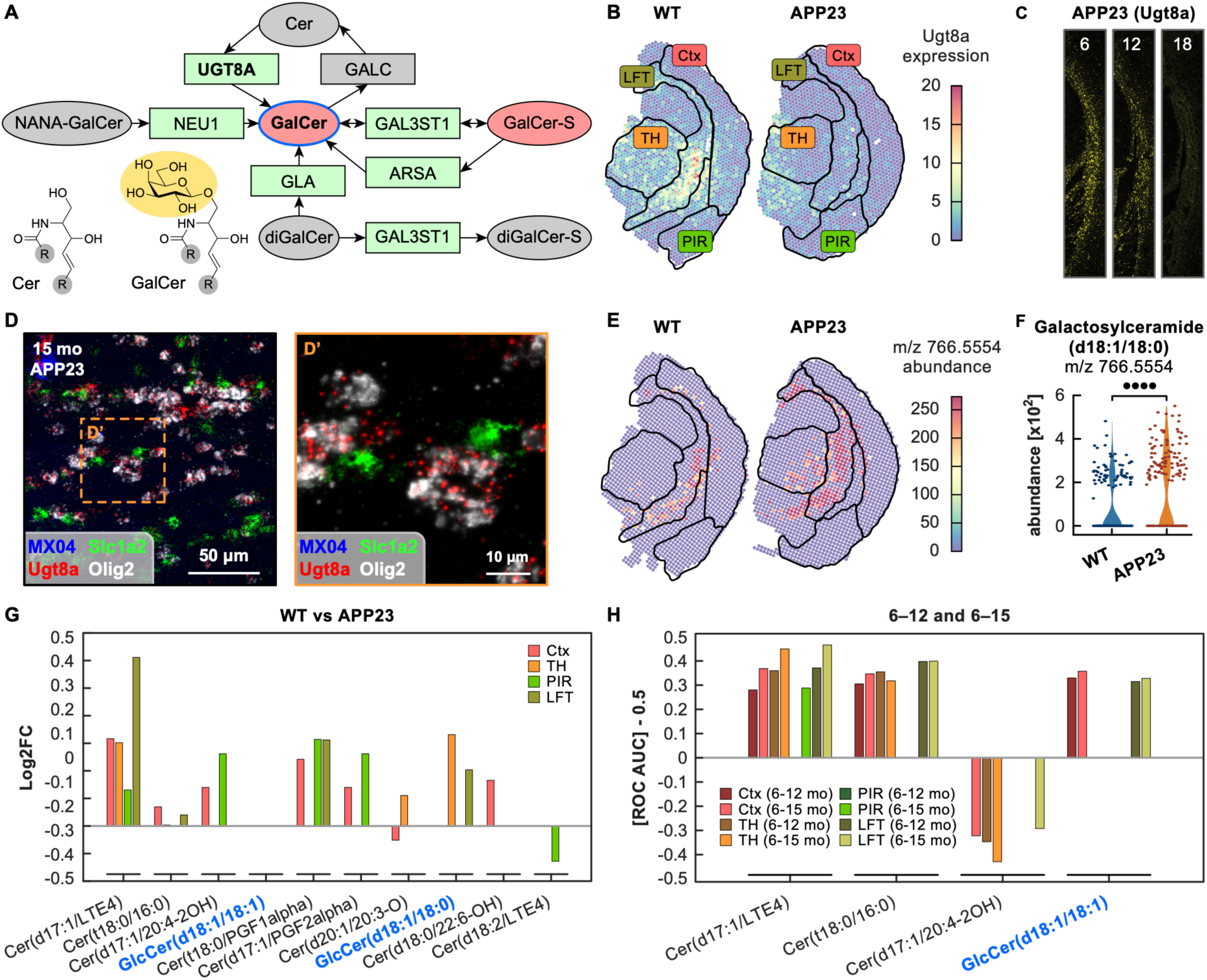
Selective remodeling of the sphingomyelin/galactosyl-ceramide in myelinated white matter tracts. **(A)** KEGG sphingomyelin pathway annotated with iMIST data. Metabolites are shown as ovals, enzymes as rectangles; red = upregulated, green = downregulated in APP23 versus WT, and gray = not significant or undetected. Galactosyl-ceramide (GalCer) is circled in blue, and the myelin-synthesizing enzyme UGT8A is highlighted in bold. (**B**) Spatial transcriptomic heatmap shows focal downregulation of Ugt8a in myelinated tracts (CC/LFT) of APP23 mice. **(C)** RNAscope validates a stepwise loss of Ugt8a transcripts from 6-to-12-to-18 months, mirroring lipid changes. **(D)** At higher magnification (63x), Ugt8a expression co-localizes with oligodendrocyte marker Olig2 (gray). Astrocyte biomarker Slc1a2 shown in green with Aβ plaque staining dye MX04 shown in blue. **(E)** Ion count maps of m/z 766.5554 (GalCer d18:1/18:0) demonstrate its selective accumulation in the LFT of APP23 brains compared with WT. **(F)** Violin plot quantifying GalCer(d18:1/18:0) signal confirms a significant increase in APP23 (Mann– Whitney U, **** P < 0.0001). **(G)** Regional changes of ceramide-related metabolites at 18 months. Bar height represents log_2_ fold change (APP23–WT) for cortex (Ctx), thalamus (TH), piriform cortex (PIR), and lateral fiber tract (LFT). GalCer(d18:1/18:0) is the sole GalCer species elevated across regions. **(H)** Age-related ROC analysis (6→12 mo and 6→18 mo) reveals age- and region-specific shifts; bars show ΔAUC–0.5 for each measured ion. GalCer(d18:1/18:1) (blue) is the only lipid significantly altered in iMIST with age in the CC/LFT.

Together, the histological anchoring (**Fig. 4C**), pathway integration (**Fig. 4B, E, F**), time-resolved metabolite profiling (**Fig. 4D**), microglial activation readouts (**Fig. 5A–C**), tentatively assigned lipid-isomer trajectories (**Fig. 5D–F** and **Fig. 6D–F**), and oligodendrocyte biosynthetic markers (**Fig. 6F–G**) provide an age-dependent intensification and expansion of both microglial and oligodendrocyte lipid-metabolic programs. These data reveal a unifying temporal mechanism in which aging progressively amplifies glial homeostatic dysfunction, deepening metabolic stress across gray and white matter through distinct, cell-type–specific pathways.

## Discussion

Growing evidence argues that AD pathogenesis may result from a gradual failure of glial lipid and energy regulation that undermines neuronal support. This concept links genetic and environmental risks— *APOEε4*, lipid transport defects, inflammation, and aging—under a shared mechanism of metabolic imbalance whose regional expression depends on cellular context. Until now, however, no approach could simultaneously map these metabolic and transcriptional disturbances within a unified spatial framework to link molecular imbalances to global brain-wide dysfunction. The iMIST platform fills this gap by integrating MALDI metabolite imaging, histology, and spatial transcriptomics in a single tissue section. This allows the spatially dependent impacts of glial stress to be detected at both the transcriptional and metabolic levels, anchored by imaging-based registration to visible pathology.

While prior work has attempted to bridge spatial metabolomics with transcriptomics, such efforts have generally been narrow in scope. For example, Vicari et al. applied MALDI-matrix directly to Visium slides in the context of analyzing Parkinson’s disease brain slices^63^, but the analysis focused on dopamine mapping and a few other small-molecule metabolites rather than a comprehensive survey of lipid chemistry. As MALDI ionization is highly dependent on surface conductivity, standard glass slides are sub-optimal conductors, which further constrains sensitivity and chemical breadth. Thus, our iMIST approach leveraging a conductive alumina membrane for matrix-free ionization for MALDI-MSI allows a seamless transition from sectioning to MSI to fluorescence imaging to RNA capture. Another key advantage of the membrane is a more homogenous ionization profile, improving the MSI analysis^64^. These qualities improve the accessibility and versatility of iMIST as a spatial multi-omic profiling method. Critically, the iMIST analytical backbone, D-iMISTer, introduces a unique consistency-validation and cross-regional concordance framework for spatial multi-omics that enables more accurate calls of perturbations among samples and spatial domains. This methodological refinement increases the transcriptomic Jaccard overlap and MS Venn diagrams statistical confidences beyond single-comparison thresholding can achieve. This analysis approach can reduce false-positive feature calls in any spatial omics study comparing multiple anatomical or pathological zones. Future studies applying iMIST to other disease models or tissue types can adopt the same validation logic without modification, underscoring the platform’s potential as a versatile framework for spatial multi-omic discovery.

From a biological angle, our work delivers the first integrated transcriptomic and metabolomic analysis of AD pathology in a rodent model system. The validity of iMIST data is evident from the overlap between our PAG module and prior A1^53^, DAM^22^ and “Purple”^51^ gene sets. The 43 newly identified PAG genes and dozens of PAM m/z species showcase the method’s discovery power as a basis for developing new hypotheses based on a wholistic view of gene expression levels and metabolite abundance across discrete brain regions. This adds to the understanding of how plaques may arise from, and then act as, focal intensifiers of global metabolic imbalance. The observed microglial lipid remodeling near plaques and oligodendrocyte lipid-synthesis imbalance in myelinated tracts—reflect a shared, age-amplified decline in glial energy and lipid regulation.

For example, we observe a pronounced rise in PA and its PC and PE precursors. This coincides with the concurrent rise of PLD4 transcript, which encodes a lysosomal enzyme supporting lipid catabolism. While named for phospholipase D activity, mammalian PLD3 and PLD4 are endolysosomal glycoproteins with minimal classical phospholipase activity. Instead, they function as 5’ exonucleases and modulators of nucleic acid sensing pathways in microglia and other immune cells, roles that can secondarily influence lipid turn over and membrane composition.^55, 65, 66^ The expression patterns of PLD4 has previously been implicated in systemic autoimmune diseases^67^ and myelin remodeling^68^. Other members of this family, such as PLD3 have been linked with AD risk and plaque-associated axonal injury.^9, 69^ The synchronous gene expression and lipid alteration identified by iMIST may suggest endolysosomal stress—which is related to lipid remodeling pathway in plaque-associated microglia—an association that bulk omics would have missed.

Equally suggestive are the white matter-restricted metabolic and gene signatures found in the CC and LFT—regions largely devoid of plaques and consisting of myelinated axon bundles. Detection of selective accumulation of GalCer species together with down-regulation of Ugt8a, might be related to an early myelin lipidopathy that is independent of age-related changes or Trem2-mediated microgliosis.^70, 71^ In contrast, this white matter-specific aberrant Ugt8a-GalCer circuit may reflect dysfunctional interactions between oligodendrocytes and the perturbed axons emanating from distal plaque-rich regions. Ugt8a encodes UDP-galactose ceramide galatosyltransferase, the enzyme required for the synthesis of galactocerebroside and sulfatides, which are the core galactolipids of CNS myelin essential for node formation, axo-oligodendrocyte interactions, and the stability of myelin sheaths.^72^ Loss of Ugt8a activity impairs these structural features, which can alter oligodendrocyte maturation, and has been linked to conduction deficits even when myelin protein expression appears intact.^73^ Because GalCer lipids are core myelin constituents, the multiomic findings implicate a model involving oligodendrocyte homeostasis before or independent of proximal plaque deposition, potentially coupling other AD risk factors such as APOE-associated risks and myelin maintenance. This notion is consistent with recent single-nuclei sequencing studies in AD patient brains^11,41,42,74^ showing detrimental oligodendrocyte signatures and perturbed myelination pathways, which were exacerbated by *APOEε4*. Based on prior work showing that galactolipid deficiency disrupts nodal organization and increases the pool of immature oligodendrocytes^9, 75^, we hypothesize that early Ugt8a downregulation in plaque-distant white matter may reflect a compensatory phenotype that eventually renders these tracts more susceptible to degeneration.

Broadly, our results illustrate the valuable lines of questioning revealed through integrated multiomics analysis that is possible through iMIST sampling. By integrating metabolite, transcript, and histological data within a single spatial framework, iMIST links local homeostatic or pathological events to systemic tissue changes, providing mechanistic insight and a generalizable platform for mapping biological processes at cellular resolution. The D-iMISTer bioinformatic pipeline also offers a generalizable framework for comparing datasets across multiple samples and tissue regions that is immediately applicable for other spatial analysis goals. We anticipate that the specific use of only commercially-available components makes iMIST broadly accessible to the research community and also suggests that the alumina membrane ionization approach might bring metabolomic measurements to other spatial nucleic acid methods.

## Resource availability

### Lead contact

Further information and requests for resources and reagents may be directed to the corresponding author Adam Abate and Carlo Condello.

### Materials availability

Materials used in this study are available upon request in accordance with the Materials Transfer Agreement (MTA) of University of California San Francisco.

## Acknowledgments

The authors gratefully acknowledge Prof. William F. DeGrado for his contributions to manuscript preparation and insightful discussions. This work was supported by grants from the National Institutes of Health (NIH) (R01NS130876 to A.R.A and C.C.; R01AI178795, R01AI149699, and R42GM143989 to A.R.A.; P01AG002132 and R01AG082141 to C.C.). This work was also supported by funding from the UCSF Sandler Program for Breakthrough Biomedical Research (A.R.A. and C.C.) and Chan Zuckerberg Biohub (A.R.A.). L.X. was supported by the National Key R&D Program of China (Grant No. 2024YFF0725700), the National Natural Science Foundation of China (Grant Nos. 32371516) and Central Guidance Fund for Local Science and Technology Development Projects (Xinjiang) (ZYYD2025QY22HZ). H.Y. was supported by the NIH (R00AG084926), BrightFocus Foundation postdoctoral fellowship (A2020039F), and the UCSF Sandler Program for Breakthrough Biomedical Research postdoctoral award (7000/7002124). J.L. was supported by the Alzheimer’s Association research fellowship (AARFD-24-1308438). We appreciate Nannan Tao from Bruker Nano Surfaces for the Bruker software support. We also appreciate Dr. Man Wang from Institute for Neurodegenerative diseases at UCSF for mass spectrometry support, and Haoran Chen from Xi’an Jiaotong University for data analysis support.

## Author contributions

L.X., H.Y., A.R.A., and C.C. designed research; L.X., H.Y., A.O., and C.C. performed experiments and collected data; L.X., H.Y., J.L., X.L., C.M., A.R.A., and C.C. analyzed data; L.X., H.Y., C.M., A.R.A., and C.C. wrote the paper; and A.R.A. and C.C. supervised the study. All authors read and approved the manuscript.

## Declaration of interests

Adam R. Abate and Linfeng Xu are named inventors on patent applications entitled “Multimode Omics of Single Tissue Sample,” including U.S. Provisional Patent Application No. 63/245,371, PCT Application No. PCT/US2022/043624 (published as WO2023043897A1), and U.S. Patent Application No. 18/690,840 (published as US20240392354A1). The applications relate to methods for integrating spatial metabolomic, transcriptomic, and imaging measurements from the same tissue section and are relevant to the methods described in this study.

## Notes

### Summary of Updates

Revision 2; Figures revised; Declaration of interest updated; Manuscript revised; Abstract revised.

