## Supplemental Information for "Spatial transcriptome-metabolome integration reveals region-specific glial lipid dysregulation linked to Alzheimer’s pathology"

Table of Contents

| Table S1. Key Resources |
| --- |
| Resource Availability |
| Lead Contact |
| Materials Availability |
| Experimental Model and Subject Details |
| Animals |
| Method Details |

### Table S1. Key Resources

| **REAGENT or RESOURCE** | **SOURCE** | **IDENTIFIER** |
| --- | --- | --- |
| **RNA scopes** | | |
| UGT8a (Mm-Ugt8a-C2) | ACDBio | Catalog number: 480611-C2 |
| PLD4 (Mm-Pld4-C2) | ACDBio | Catalog number: 1222001-C2 |
| CSF1R (Mm-Csf1r-C2) | ACDBio | Catalog number: 428191-C2 |
| CLEC7A (Mm-Clec7a-C1) | ACDBio | Catalog number: 532061 |
| SLC1A2 (Mm-Slc1a2-C1) | ACDBio | Catalog number: 441341 |
| OLIG2 (Mm-Olig2-C2) | ACDBio | Catalog number: 447091-C2 |
| **Biological Samples** | | |
| B6-Tg(Thy1-HuAPP[751]Sw)23Apoe^tm1.1(APOE*4)Adiuj/J | Hunters point (UCSF) | Animal IDs: DD330, EE14319,  EE16522, DD15478, ER11465 |
| B6-Tg(Thy1-HuAPP[751]Sw)23Apoe^tm1.1(APOE*4)Adiuj/J  ApoE-KI TM/TM,  HuAPP qPCR - | Hunters point (UCSF) | Animal IDs: DD328 |
| B6-Tg(Thy1-HuAPP[751]Sw)23HOZ | Hunters point (UCSF) | Animal IDs: FF2957 (15 mo) |
| B6 | Hunters point (UCSF) | Animal IDs: FF20766 (6 mo) FF23783 (15 mo) |
| **Chemicals, Peptides, and Recombinant Proteins** | | |
| LDS722 | Exciton | Cat. No. 07220 |
| FluoroMyelin™ Green Fluorescent | Invitrogen | Cat. No. F34651 |
| DIUTHAME MALDI membrane | Hamamatsu | Cat. No. A13331-5019-1 |
| PermaFluor mounting medium | Epredia | Cat. No. TA-030-FM |
| L-Arginine | Sigma-Aldrich | Cat. No.11009 |
| **Critical Commercial Assays** | | |
| NextSeq2000 |  | Illumina |
| **Deposited Data and code** | | |
| Sequencing & MALDI imaging raw and processed data, code, supplementary figures, data, and tables | Dryad | DOI: 10.5061/dryad.m37pvmd9h |
| **Software and Algorithms** | | |
| Seurat | Seurat | V3.2 |
| MATLAB | MathWorks | R2021b |
| R code | The R Project | V4.1.3 |
| ftmsControl | Bruker | V2.3.0 |
| flexImaging | Bruker | V5 |
| SCiLS Lab | Bruker | V2023b |
| ImageJ | Fiji | V1.53 |
| MetaboAnalyst | MetaboAnalyst | V6.0 |

### Resource Availability

**Lead Contact**

Further information and requests for resources and reagents may be directed to the corresponding author Adam Abate and Carlo Condello.

**Materials Availability**

Materials used in this study are available upon request in accordance with the Materials Transfer Agreement (MTA) of University of California San Francisco.

### Experimental Model and Subject Details

### Sectioning → membrane placement → MALDI → histology → Visium → sequencing.

**Animals**

All comparisons were conducted on brains from sex-matched and liter-mate animals. Animals were maintained in a facility accredited by the Association for Assessment and Accreditation of Laboratory Animal Care International in accordance with the Guide for the Care and Use of Laboratory Animals. All procedures for animal use were approved by the University of California, San Francisco’s Institutional Animal Care and Use Committee. Tg(APP23) mice, which express human APP (751-aa isoform) containing the Swedish mutation under the control of the Thy-1.2 promoter, were maintained on a C57BL/6 background.^[1]^ In this study, Tg(APP23) female mice were 6-, 12-, 15-, and 18-months old and wildtype 6-, 12-, 15-, and 18-month-old liter-mate as controls. Mouse brain samples were harvested, flash frozen, and cryo-sectioned following standard procedures. Flash frozen mouse brains were coronal cryo-sectioned (14-μm thickness) and mounted on 10x Visium Spatial Gene Expression slide for spatial transcriptomics. Right after mounting, DIUTHAME MALDI membrane^[2]^ was placed on top of the brain section before the tissue is dried. A dry ice was placed right under the slide to keep the tissue frozen.

**MALDI imaging**

The membrane was brought to room temperature and then the membrane on sample slide was sprayed with MeOH using HTX TM-Sprayer (0.125 ml/min, 30 °C, HH pattern, 1 pass) to absorb the metabolites from the brain. MALDI imaging was performed in Bruker SolariX T7. The ftmsControl (Bruker Daltonics, v 2.3.0) and the flexImaging (Bruker Daltonics, v 5.0) software were employed for data acquisition. The mass calibration was performed with L-Arginine (50% MeCN in H_2_O with 0.1% formic acid) in the ESI-MS positive-ion mode. The slide was scanned with an Epson scanner (highest resolution setting) and then loaded onto SolariX using Bruker MTP slide adapter II. MALDI imaging was performed in positive-ion mode at the mass range of m/z 150–3000 with “small” laser focus setting (80-μm beam size). For each MALDI experiment, 200 laser shots were accumulated with a 100-μm lateral resolution.

**Aβ plaque staining and histology.**

The post-MALDI sample slide was put in -20 °C MeOH solution containing 0.05% LDS722 (w/v) for 20 min, which the solution was then brought to room temperature, and then further dried under vacuum. The brain sections were imaged with Leica Dmi8 epi-fluorescence microscope equipped with a 20x HC PL FLUOTAR L dry lens (0.4 NA), red and green fluorescence filters (630 nm and 527 nm emission wavelength) with 5 mS exposure time, a DFX9000GT camera at 1 x zoom and 2048x2048-pixel resolution. The Leica LAS X Navigator module within the Leica LAS X software was utilized to achieve full brain histology. For each field-of-view experiment, the optical plane was auto-focused on the highest sensitivity setting. Due to MALDI membrane blocking the path of the light, the slide was loaded upside down to image through the glass slide. Brain micrographs were acquired and merged using the Leica LAS X Navigator module in Leica LAS X software to stitch full brain histology. The image was then brought to Fiji software which was used to segment the Aβ plaques and cerebral amyloid angiopathy (CAA) in the brain, determined by morphology and staining patterns.

**Spatial gene expression.**

Standard 10x Visium Spatial Gene Expression was followed with an updated protocol of using proteinase K instead of pepsin for tissue permeabilization step as we found proteinase K can dissolve the tissue more efficiently which allowed the removal of tissue bound MALDI membrane for downstream RNA capture. The proteinase K lysis buffer was prepared with 0.94 mg/ml proteinase K (NEB) dissolved in ddH_2_O of 92.81 mM Tris-HCl pH 8.0, 0.94% NP-40 and 9.38 mM EDTA. Spatial gene expression libraries were generated following 10X Genomics Visium Gene Expression and Tissue Optimization protocols according to the manufacturer’s recommendations (Visium Spatial Gene Expression Reagent Kits—Tissue Optimization User Guide, document CG000238 Rev E, 10X Genomics, (February 2022); Visium Spatial Gene Expression Reagent Kits—User Guide, document CG000239 Rev F, 10X Genomics, (January 2022) and Methanol Fixation, H&E Staining and Imaging for Visium Spatial Protocols, document CG000160 Rev C, 10X Genomics). Libraries were sequenced using a NextSeq2000 sequencing system (Illumina). The length of read 1 was 28 bp, while the length of read 2 was 150 bp long.

**MALDI-MSI data processing.**

The data was visualized by FlexImaging and analyzed with SCiLS lab (Bremen, Germany). The raw data were imported into the SCiLS lab, and spectra were normalized against the total ion count. Subsequently, the SCiLS Lab Pro (Bruker Daltonics, v 2023b) and the SCiLS Lab API (Bruker Daltonics, v 6.0) software were used to export the data as imzML format which was then used for R package Cardinal MSI (Cardinal 3.0) software. Because the high-resolution massspect shift can occur during the data collection, and to ensure m/zs across samples match, we binned m/z peaks to a reference peak list. The bin size was ±2.5 ppm. Then the Cardinal MSI software was used to convert the imported imzML data to a csvfile spreadsheet for downstream multiomics analysis. For kinetics studies, ROC analyses were performed using SCiLS Lab Pro (Bruker Daltonics, v 2023b) software to track abundance changes in m/z peaks over time.

**Metabolite identification.**

All metabolites were assigned by subjecting the m/z peaks to human metabolome database (HMDB)^[3]^. The m/z peaks were queried in the positive-ion mode with adduct types of [M+H], [M+H-H2O], [M+Na], [M+K] with molecular weight tolerance of 10 ppm. All metabolites were downloaded and put into an array per m/z peak for downstream analysis. Using the KEGG ID of each tentatively assigned metabolites, integrated metabolome and transcriptome KEGG pathway analysis was performed using MetaboAnalyst 6.0^[4]^. The list of tentatively assigned metabolites for identified m/z peaks are available as a supplementary data (Supplementary Data S3).

**Visium data processing.**

Sequenced libraries were mapped to a custom genome using Space Ranger (1.3.1; 10X Genomics)^[5]^. A custom mm10 mouse reference genome was generated by adding hFAD plasmid as an extra chromosome following 10x recommendations. The 1–849 bp of the Thy1 sequence was removed from the original plasmid sequence after observing an overlap with an adjacent gene following a blast similarity analysis. General mm10 reference and the corresponding gene transfer format (GTF) files were downloaded from Ensembl (https://useast.ensembl.org/index.html). Gene body coverage analysis was performed using the target sequences from the possorted_genome_bam.bam files using Samtools (https://www.htslib.org/). Analyses of sequencing saturation and median genes per spot as functions of mean reads per spot were performed using Space Ranger and R package Seurat.

**Multiomics data analysis.**

The D-iMISTer analytical framework is designed around two core principles: (1) all data modalities must originate from the same tissue section to eliminate inter-section alignment errors, and (2) differential features must be validated across spatially related regions to distinguish biological signals from regional noise. The first principle is achieved experimentally through the iMIST pipeline, which sequentially acquires MALDI-MSI, histology, and spatial transcriptomics from a single cryosection. The second principle is implemented computationally through a consistency-supported cross-validation filter that requires each differentially expressed feature to show concordant fold-change direction across partner comparisons. Together, these principles ensure that the spatial multi-omic signatures reported in this study are both technically precise and biologically reproducible.

**Data processing of spatial transcriptomics**

The spatial transcriptomics data obtained with Visium were processed and analyzed using R (v4.1.3), the single-cell genomics toolkit Seurat^[6]^. The Aβ plaque staining images were transformed to binary mode and the plaque were manually selected based on morphology using the software Fiji (v1.53)^[7]^. Then the images were annotated based on tissue morphology using the interactive application Loupe Browser (10X Genomics, v6.3.0). The filtered count matrices obtained from spaceranger were used for subsequent analysis. To the filtered count matrices, we performed additional filters using Loupe Browser. In particular, spots without the tissue, spots including more than 38% mitochondrial genes or less than 50 unique genes were removed. With the filtered gene counts, the data were normalized and subjected to a basic analytical workflow using functions from the Seurat R package. The *SCTransform* function was used for normalization and variance stabilization and was followed by dimensionality reduction via PCA (*RunPCA*). Data were integrated with the *RunHarmony* function from the harmony R package^[8]^ using group.by.vars = ‘Sample.ID’ (which indicates the sample of origin), assay.use = ‘SCT’ and reduction = ‘pca’ as parameters. A shared nearest neighbor graph was constructed based on the first 30 principal components (*FindNeighbors*). Finally, a uniform manifold approximation and projection (UMAP) embedding was computed based on the first 30 principal components (*RunUMAP*); this was followed by graph-based clustering (*FindClusters*). Marker genes for each identified cluster were calculated using the function *FindAllMarkers* with default parameters, while a nonparametric Wilcoxon rank-sum test and the Bonferroni correction were used for P-value adjustment. Only genes with adjusted P-value lower than 0.05 were considered differentially expressed.

**Data processing of mass spectrometry imaging**

Next, MSI and corresponding Aβ plaque staining images were aligned using the Image processing package by MATLAB (Table S1). To be more specific, the MSI data first underwent data selection of lipid Region from 738 to 938 m/z to identify and remove data points that were located outside of the tissue sections. Then the MSI and corresponding Aβ plaque staining image were processed to binary image by edge detection function to show the boundary of the tissue sections clearly. After that, the binary image of MSI was rotated, scaled and aligned to the binary image of corresponding Aβ plaque staining image. Finally, following the 10x format, the location of each MSI spot on the Aβ plaque staining image was generated into a csv file.

**Calculation of plaque area in each RNA spot and MSI spot**

The plaque area within each RNA and MSI spot was calculated. Using the geometric centers of each RNA spot as determined from the tissue_positions_list.csv output from Spaceranger, we drew circles with 50 µm diameter. Within each circle, the number of Aβ plaque pixels were counted on the corresponding Aβ plaque staining image and count the number of plaque pixels within each circle. To characterize the surrounding plaque information for each RNA spot, we also draw circles with a diameter of 250 µm around each spot and count the plaque pixels inside these larger circles. This method allows us to add plaque information to each RNA spot. Similarly, we apply this approach to each MSI spot, incorporating plaque information in the same manner.

**WGCNA analysis of plaque induced genes**

With the filtered RNA spots, we chose genes upregulated in APP23 for WGCNA analysis^[9]^. The WGCNA was performed using the modified pyWGCNA function from the omicverse package in python. This resulted in 13 modules, and for each module, we calculated the mean gene log2(FC), which is log2 average fold change of gene expression level between APP23 and WT. Then the GLM (generalized linear model) function from SciPy was used to fit Aβ plaque numbers and gene expression level based on APP23. From that, the mean Aβ plaque log2(FC) for the corresponding genes was calculated. Finally, the scatter plot was generated between the mean gene log2(FC) and the mean plaque log2(FC)^[10]^. The module 10 (M10) stood out which was both high in gene expression level change and Aβ plaque numbers between APP23 and WT. A venn diagram was finally generated to show the overlap among the genes from M10, DAM^[11]^, A1^[12]^ and PIGs^10^.

To optimize cross-comparison reproducibility of differentially expressed features across brain regions, we applied a consistency-supported filtering strategy for Venn diagram generation. Rather than classifying significant features independently in each comparison, we required each feature to pass the primary significance threshold (|avg_log2FC| > 0.25 and adjusted p-value < 0.05) and additionally show the same fold-change direction in the partner comparison (same-direction |avg_log2FC| > 0.25). Additional pct filters were not used (high_pct_thresh = 0 and pct_diff_thresh = 0). This filter removes features that are significant in only one comparison but do not move consistently when Region C is compared with both Region A and Region B, thereby reducing false-positive overlap calls. Application of this strategy substantially improved Jaccard overlap across all datasets: RNA up-regulated features improved from 34.7% to 64.0%, RNA down-regulated from 20.4% to 60.5%, MS up-regulated from 49.1% to 89.9%, and MS down-regulated from 50.3% to 83.5%. The optimized Venn diagrams and corresponding CSV files are available in **Supplementary Fig. S5** and **Supplementary Data S5**.

In simpler terms, this filtering strategy addresses a basic but critical question: when we compare diseased brain regions with healthy controls, how do we know that a gene or metabolite we call "significant" is truly part of a coherent biological response rather than a region-specific noise artifact? The conventional approach counts any feature that crosses the significance threshold in a single comparison, which is analogous to accepting a scientific finding without independent replication. Our consistency filter instead demands that the feature behave consistently across multiple comparisons, mirroring the replication principle that underlies robust experimental design. This is particularly important in spatial omics because neighboring brain regions share similar cellular compositions and pathological environments; a genuine biological change should therefore propagate with the same directional trend across connected regions, whereas a technical artifact is more likely to appear in isolation.

**Annotation between spatial transcriptomics and mass spectrometry imaging**

As both of RNA spots and MSI spots are aligned to the same corresponding plaque staining images, the MSI data are aligned to RNA data. Then we identified pairs of nearest neighbors across the two datasets by checking the overlap of the geometry center of spots from each data set. As the density of RNA spots and MSI spots are similar, only if the geometry center of the MSI spot fell in the range of RNA spot, the MSI data was kept for each RNA data point, thus generating a list of MSI-RNA data point pairs. In cases where the multiple RNA data points shared the same MSI spot, this neighbor was reused to ensure a one-to-one mapping. Once the MSI-RNA data point pairs had been identified, we used these pairs to subset the raw data and produce new Seurat objects with the two aligned data modalities stored in separate assays.

**Differential Gene Expression (DGE)**

The differential expression analysis was conducted on RNA data for each RNA UMAP cluster between APP23 and WT. Then we used matplotlib package in python to generate the volcano plots. The threshold of P-value and log2 fold change were set to 0.01 and 0.2 to identify significant genes. Subsequently, the violinplot function in seaborn package was used to generate the violin plots, with hue = ‘sample’ and x = ‘RNA_UMAP_cluster’ to group by the RNA UMAP cluster. A t-test was then performed for each group to determine the significance of difference of genes. For cross-regional validation, a consistency-supported filtering strategy was additionally applied: significant genes were required to show concordant fold-change direction in partner comparisons (same-direction |log2FC| > 0.25) to be included in Venn diagram overlap analyses.

This cross-regional validation step is conceptually distinct from standard multiple-testing correction. While methods such as Benjamini-Hochberg FDR control reduce false positives within a single comparison by adjusting p-values, they do not assess whether a feature behaves consistently across independent but related comparisons. The consistency filter adds this second layer of scrutiny by treating fold-change concordance as an independent line of evidence. In practice, a gene that is upregulated in Region C versus A but downregulated in Region C versus B may pass the FDR threshold in one comparison yet fail the consistency filter, correctly flagging it as potentially artifactual. This dual-threshold approach thus provides stronger protection against spatial noise than either significance testing or fold-change filtering alone.

**Differential Metabolite Expression (DME) analyses**

In a similar way to DGE, the differential expression analysis was conducted on MS data for each RNA UMAP cluster between APP23 and WT. The MS data was normalized by Total Ion Count (TIC) method and then matplotlib package in python was used to generate the volcano plots. The threshold of P-value and log2 fold change were set to 0.01 and 0.2 to identify significant m/z peaks. We used violinplot function in seaborn package to generate the violin plots, with hue = ‘sample’ and x = ‘RNA_UMAP_cluster’ to group by the RNA UMAP cluster. A t-test was then performed for each group to determine the significance of difference of m/z peaks. Analogous to DGE, MS features passing the primary significance threshold were further filtered by requiring concordant fold-change direction in partner comparisons (same-direction |log2FC| > 0.25) for inclusion in cross-regional Venn overlap analyses.

Because MALDI-MSI data are subject to additional sources of spatial variability, including ionization efficiency gradients and matrix crystallization heterogeneity, the consistency filter is even more critical for metabolite features than for gene-expression features. By enforcing that an m/z peak must show the same directional change across multiple regional comparisons, we effectively separate metabolites that respond coherently to disease pathology from those whose abundance fluctuates due to technical variation. The improvement in Jaccard overlap metrics after applying this filter, particularly the jump from 49.1% to 89.9% for MS up-regulated features, quantitatively demonstrates that cross-regional consistency is a powerful discriminator of true biological signal in spatial metabolomics.

**Spatial display of target genes and m/z peaks**

To exhibit spatial distribution of RNA and MS, we processed APP23 and WT samples separately. Specifically, the scanpy package in python was used to normalize the total gene count to 10,000 reads per spot and apply a log_2_ transformation to the data. We calculated overall mean and standard deviation for two samples, then the expression level of each gene or m/z was clipped to the 3σ range to mitigate the impact of outliers and normalize the expression values of two samples onto a unified scale. For MS spots, only those that intersect with RNA spots were selected and applied above process.

**Spatial correlation analysis.**

The plaque values were log_2_-transformed and then normalized using min-max scaling. The Pearson correlation coefficients between Aβ plaque and each gene or m/z were then calculated for three brain regions: Ctx, PIR (Ctxsp) and TH from APP. We filtered out the genes or m/z peaks that appeared in less than 2% spots of each brain region. For MS data, m/z below 400 were excluded. For each of Ctx, PIR (Ctxsp) and TH clusters, 10 genes or m/z values with the highest and the lowest correlations were chosen. Finally, a correlation matrix was constructed and a heatmap was generated using heatmap function of seaborn package as shown in Figure 3.

**Integrative pathway analysis of transcriptomics and metabolomics.**

Statistical meta-analysis module in MetaboAnalyst 6.0^[13]^ was used to perform integrative analysis of transcriptomics and metabolomics data at the KEGG pathway level. The identified metabolites and genes (p-value < 0.05) from differential gene expression and differential metabolite abundance analyses were queried. The corresponding names (genes and KEGG compound ID) with their fold changes were used as an input. To assess all the identified metabolites and genes, we multiplied the fold change with the log_2_ of significance. For pathway analysis, integrated metabolic pathways was selected with hypergeometric test for enrichment analysis.

**RNA Scope.**

RNA scope experiments were followed based on the Multiplex Fluorescent Reagent Kit v2 Assay (#323100). The slides were placed under a UV lamp for approximately 24 h, and reagents such as 1x Wash Buffer and 20x SSC are prepared by warming, diluting, adjusting pH, and sterilizing. The FFPE brain sections were incubated at 60°C for 1 h, followed by deparaffinization with xylene and ethanol. RNAscope Hydrogen Peroxide was applied, followed by washing and target retrieval using a steamer, protease treatment, and hybridization of RNA probes at 40°C for 2 h. The slides were washed, and placed in the HybEZ oven pre-warmed to 40°C. Sequential hybridization of either Pld4 (Mm-Pld4-C2)or Ugt8a (Mm-Ugt8a-C2) is performed with incubation and washing steps. Opal dyes are prepared and applied to develop signals for corresponding HRP-channel probes, followed by HRP blocker. Finally, slides are counterstained with methoxy-04 (MX04) for Aβ plaque staining, followed by mounting with a coverslip with PermaFlour, and dried overnight in the dark. The prepared slides are stored at 4 °C. The brain sections were imaged with two data collection methods. (1) Leica SP8 confocal microscope using a 20x water immersion lens (1.1 NA), a white light and 405 nm lasers, a HyD detector at 512×512-pixel resolution at 1x zoom. The Leica LAS X Navigator module within the Leica LAS X software was utilized to achieve comprehensive brain histology. For each field-of-view experiment, the optical plane was auto-focused on the highest sensitivity setting. (2) high-magnification images were acquired using a 63× oil immersion objective (1.4 NA) on the Leica SP8 confocal microscope operated in sequential scan mode to prevent channel crosstalk. The following channels were used depending on probe fluorophores: 405 nm, 520 nm, 570 nm, and 690 nm. Images were collected at either 2048 × 2048 pixels (zoom 1) or 1024 × 1024 pixels (zoom 3) resolution. Z-stacks were acquired where indicated to capture three-dimensional signal distribution within the tissue section

1. Sturchler-Pierrat C, Abramowski D, Duke M, Wiederhold KH, Mistl C, Rothacher S, Ledermann B, Bürki K, Frey P, Paganetti PA, Waridel C, Calhoun ME, Jucker M, Probst A, Staufenbiel M, Sommer B. Two amyloid precursor protein transgenic mouse models with Alzheimer disease-like pathology. Proc Natl Acad Sci U S A. 1997 Nov 25;94(24):13287-92. doi: 10.1073/pnas.94.24.13287. PMID: 9371838; PMCID: PMC24301.
2. Kuwata K, Itou K, Kotani M, Ohmura T, Naito Y. DIUTHAME enables matrix-free mass spectrometry imaging of frozen tissue sections. Rapid Commun Mass Spectrom. 2020 May 15;34(9):e8729. doi: 10.1002/rcm.8729. PMID: 31951673.
3. Wishart DS, Guo A, Oler E, Wang F, Anjum A, Peters H, Dizon R, Sayeeda Z, Tian S, Lee BL, Berjanskii M, Mah R, Yamamoto M, Jovel J, Torres-Calzada C, Hiebert-Giesbrecht M, Lui VW, Varshavi D, Varshavi D, Allen D, Arndt D, Khetarpal N, Sivakumaran A, Harford K, Sanford S, Yee K, Cao X, Budinski Z, Liigand J, Zhang L, Zheng J, Mandal R, Karu N, Dambrova M, Schiöth HB, Greiner R, Gautam V. HMDB 5.0: the Human Metabolome Database for 2022. Nucleic Acids Res. 2022 Jan 7;50(D1):D622-D631. doi: 10.1093/nar/gkab1062. PMID: 34986597; PMCID: PMC8728138.
4. Xia J, Psychogios N, Young N, Wishart DS. MetaboAnalyst: a web server for metabolomic data analysis and interpretation. Nucleic Acids Res. 2009 Jul;37(Web Server issue):W652-60. doi: 10.1093/nar/gkp356. Epub 2009 May 8. PMID: 19429898; PMCID: PMC2703878.
5. Space Ranger (1.3.1; 10X Genomics)
6. Hao Y, Stuart T, Kowalski MH, Choudhary S, Hoffman P, Hartman A, Srivastava A, Molla G, Madad S, Fernandez-Granda C, Satija R. Dictionary learning for integrative, multimodal and scalable single-cell analysis. Nat Biotechnol. 2024 Feb;42(2):293-304. doi: 10.1038/s41587-023-01767-y. Epub 2023 May 25. PMID: 37231261; PMCID: PMC10928517.
7. Schindelin J, Arganda-Carreras I, Frise E, Kaynig V, Longair M, Pietzsch T, Preibisch S, Rueden C, Saalfeld S, Schmid B, Tinevez JY, White DJ, Hartenstein V, Eliceiri K, Tomancak P, Cardona A. Fiji: an open-source platform for biological-image analysis. Nat Methods. 2012 Jun 28;9(7):676-82. doi: 10.1038/nmeth.2019. PMID: 22743772; PMCID: PMC3855844.
8. Korsunsky I, Millard N, Fan J, Slowikowski K, Zhang F, Wei K, Baglaenko Y, Brenner M, Loh PR, Raychaudhuri S. Fast, sensitive and accurate integration of single-cell data with Harmony. Nat Methods. 2019 Dec;16(12):1289-1296. doi: 10.1038/s41592-019-0619-0. Epub 2019 Nov 18. PMID: 31740819; PMCID: PMC6884693.
9. Langfelder P, Horvath S. WGCNA: an R package for weighted correlation network analysis. BMC Bioinformatics. 2008 Dec 29;9:559. doi: 10.1186/1471-2105-9-559. PMID: 19114008; PMCID: PMC2631488.
10. Chen WT, Lu A, Craessaerts K, Pavie B, Sala Frigerio C, Corthout N, Qian X, Laláková J, Kühnemund M, Voytyuk I, Wolfs L, Mancuso R, Salta E, Balusu S, Snellinx A, Munck S, Jurek A, Fernandez Navarro J, Saido TC, Huitinga I, Lundeberg J, Fiers M, De Strooper B. Spatial Transcriptomics and In Situ Sequencing to Study Alzheimer's Disease. Cell. 2020 Aug 20;182(4):976-991.e19. doi: 10.1016/j.cell.2020.06.038. Epub 2020 Jul 22. PMID: 32702314.
11. Keren-Shaul H, Spinrad A, Weiner A, Matcovitch-Natan O, Dvir-Szternfeld R, Ulland TK, David E, Baruch K, Lara-Astaiso D, Toth B, Itzkovitz S, Colonna M, Schwartz M, Amit I. A Unique Microglia Type Associated with Restricting Development of Alzheimer's Disease. Cell. 2017 Jun 15;169(7):1276-1290.e17. doi: 10.1016/j.cell.2017.05.018. Epub 2017 Jun 8. PMID: 28602351.
12. Liddelow SA, Guttenplan KA, Clarke LE, Bennett FC, Bohlen CJ, Schirmer L, Bennett ML, Münch AE, Chung WS, Peterson TC, Wilton DK, Frouin A, Napier BA, Panicker N, Kumar M, Buckwalter MS, Rowitch DH, Dawson VL, Dawson TM, Stevens B, Barres BA. Neurotoxic reactive astrocytes are induced by activated microglia. Nature. 2017 Jan 26;541(7638):481-487. doi: 10.1038/nature21029. Epub 2017 Jan 18. PMID: 28099414; PMCID: PMC5404890.
13. Xia J, Psychogios N, Young N, Wishart DS. MetaboAnalyst: a web server for metabolomic data analysis and interpretation. Nucleic Acids Res. 2009 Jul;37(Web Server issue):W652-60. doi: 10.1093/nar/gkp356. Epub 2009 May 8. PMID: 19429898; PMCID: PMC2703878.
