## Supplemental Figures for "Spatial transcriptome-metabolome integration reveals region-specific glial lipid dysregulation linked to Alzheimer’s pathology"

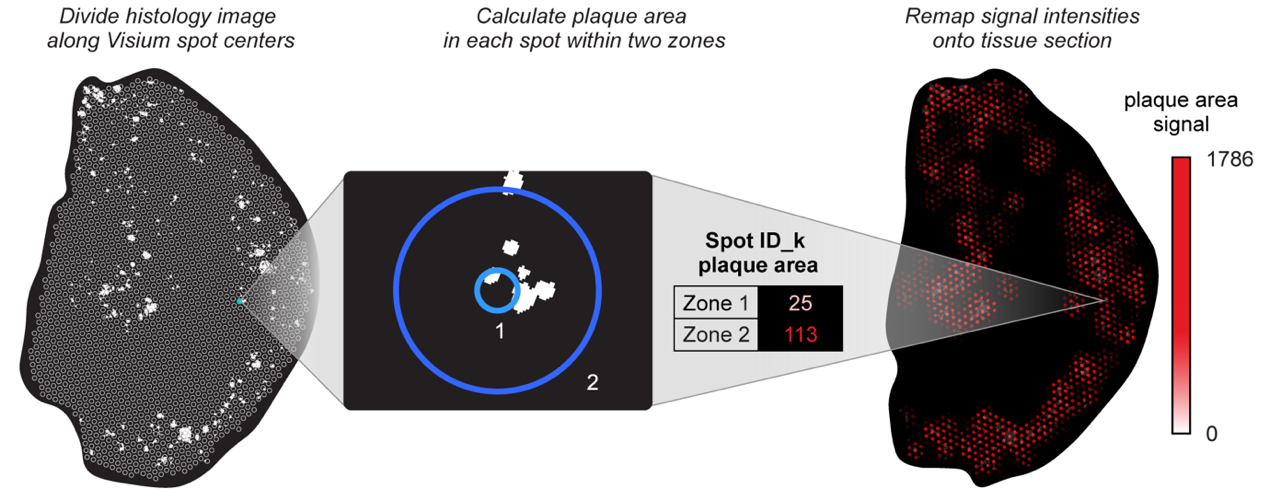

**Supplementary Figure S1. Definition of plaque area within each spatial spot.** For every spot, two plaque-area values were computed by summing the area of plaque pixels from within two concentric circular zones, centered and indexed on each Visium spot: Zone 1 (D = 50-μm) and Zone 2 (D = 250-μm).

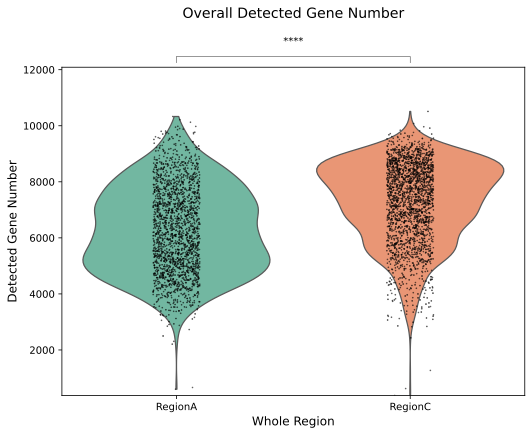

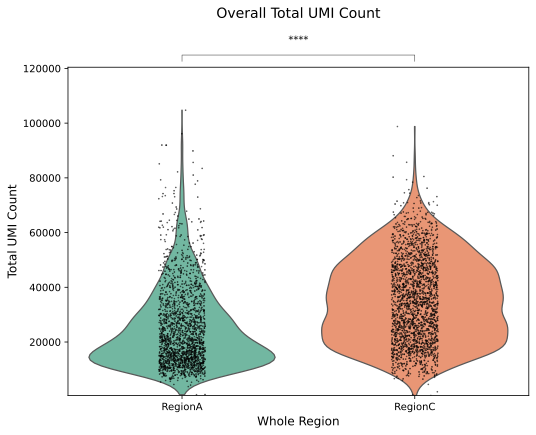

**Supplementary Fig S2. Quality metrics for 10x Visium spatial transcriptomics data.** Violin plots showing the distribution of (left) the number of detected genes per spot and (right) the total UMI counts per spot for the Visium datasets (WT in green [RegionA] and APP23 in orange [RegionC]). Each violin depicts the per-spot variability across the tissue section, highlighting overall sequencing depth and capture efficiency for each sample.

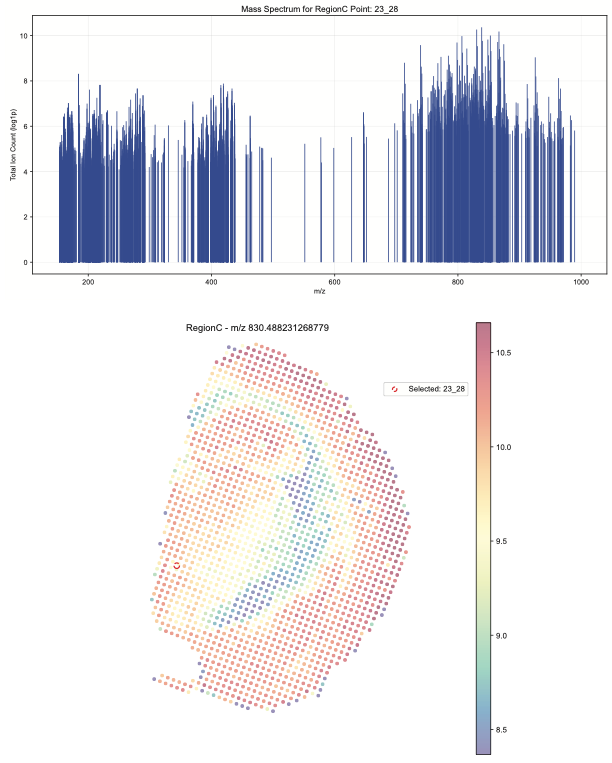

**Supplementary Fig S3. Conductive-membrane–assisted MALDI-MSI efficiently ionizes within the lipid mass-range.** A representative positive m/z ion mass spectrum collected from a mouse brain section mounted on a 10x Visium slide and overlaid with the conductive alumina membrane shows robust peak density and signal-to-noise across m/z 100–1000. Spectra were acquired on a Bruker Solarix 7.0T with a 100-µm raster and total ion count normalized and binned. The observed mass range encompasses abundant lipid species (e.g., glycerophospholipids and sphingolipids), demonstrating that the membrane enables sensitive, matrix-free MALDI-MSI suitable for lipidomics within iMIST. Spot 23_28 selected and its corresponding mass spectrum shown above.

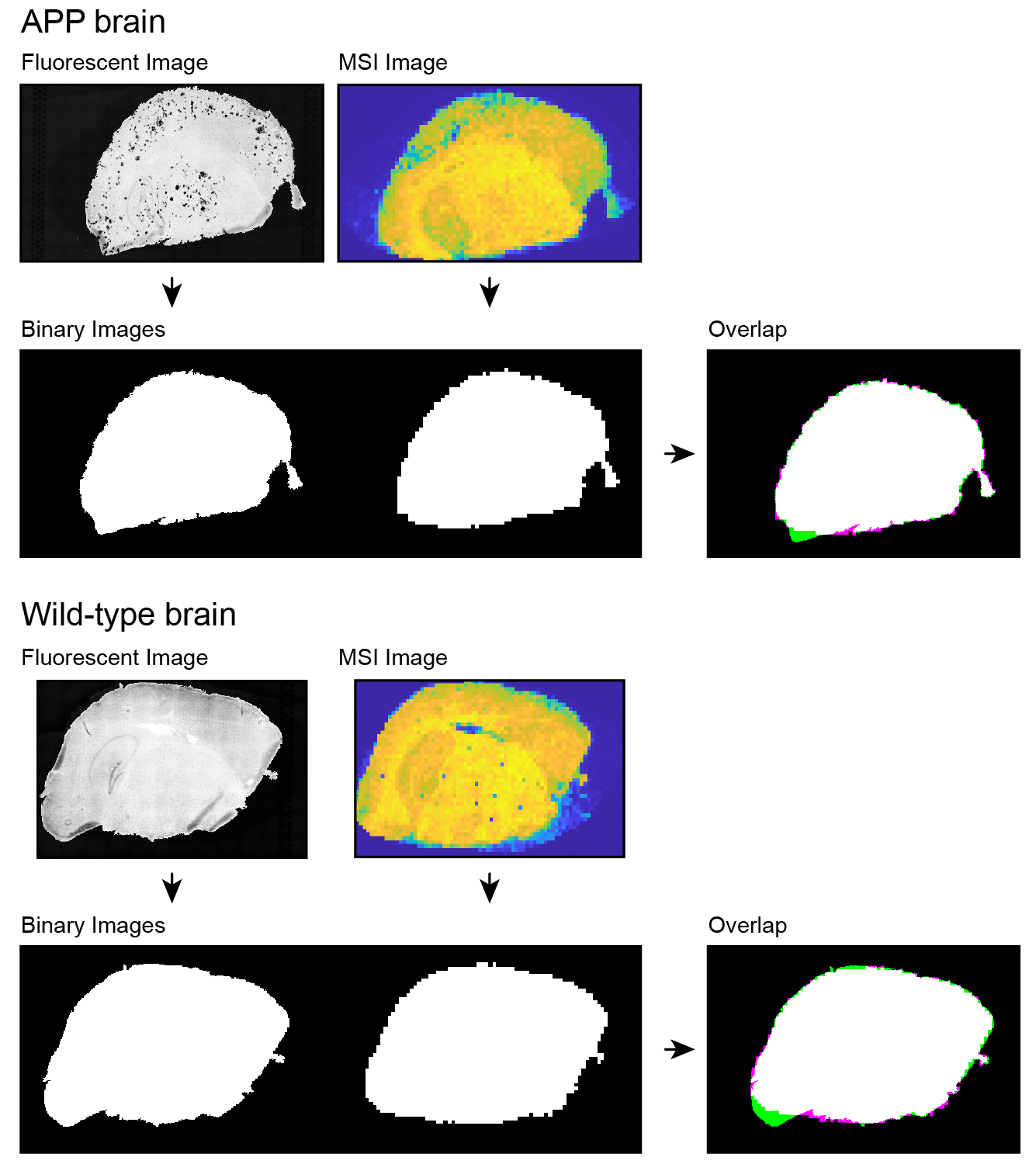

**Supplementary Fig S4. Analysis of lateral metabolite diffusion.** Overlay testing from APP brain (*top set*) and wild-type brain (*bottom set*). In each set: left image shows fluorescent staining of brain, right image shows MSI output for total m/z signal, with bottom images showing the binary image extractions and the composite overlay. Limited misregistration shows that overall diffusion of metabolites was not significant.

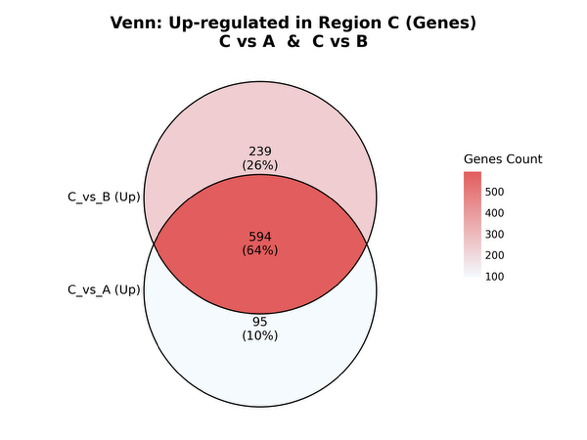

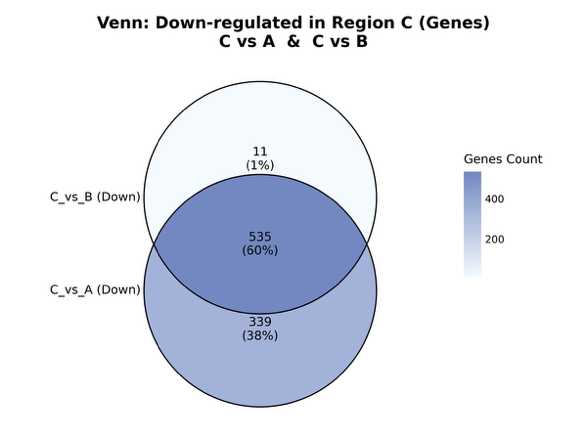

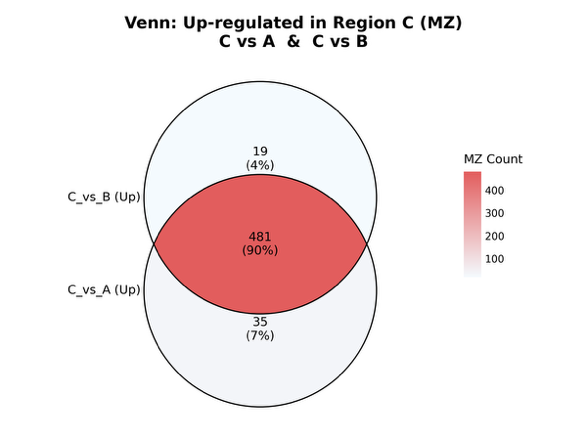

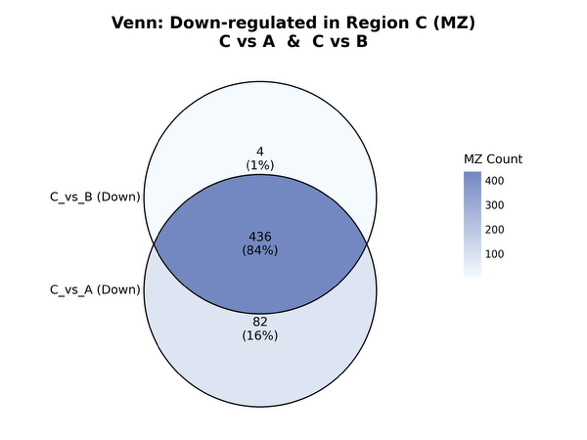

**Supplementary Fig S5. Venn Diagrams to show the reproducibility of iMIST.**

*Left*, up regulated genes and m/z peaks when APP brain sections compared to two wild-type brain sections; *Right*, down regulated genes and m/z peaks when APP brain section compared to two wild-type brain sections. Based on Supplementary Data S5.

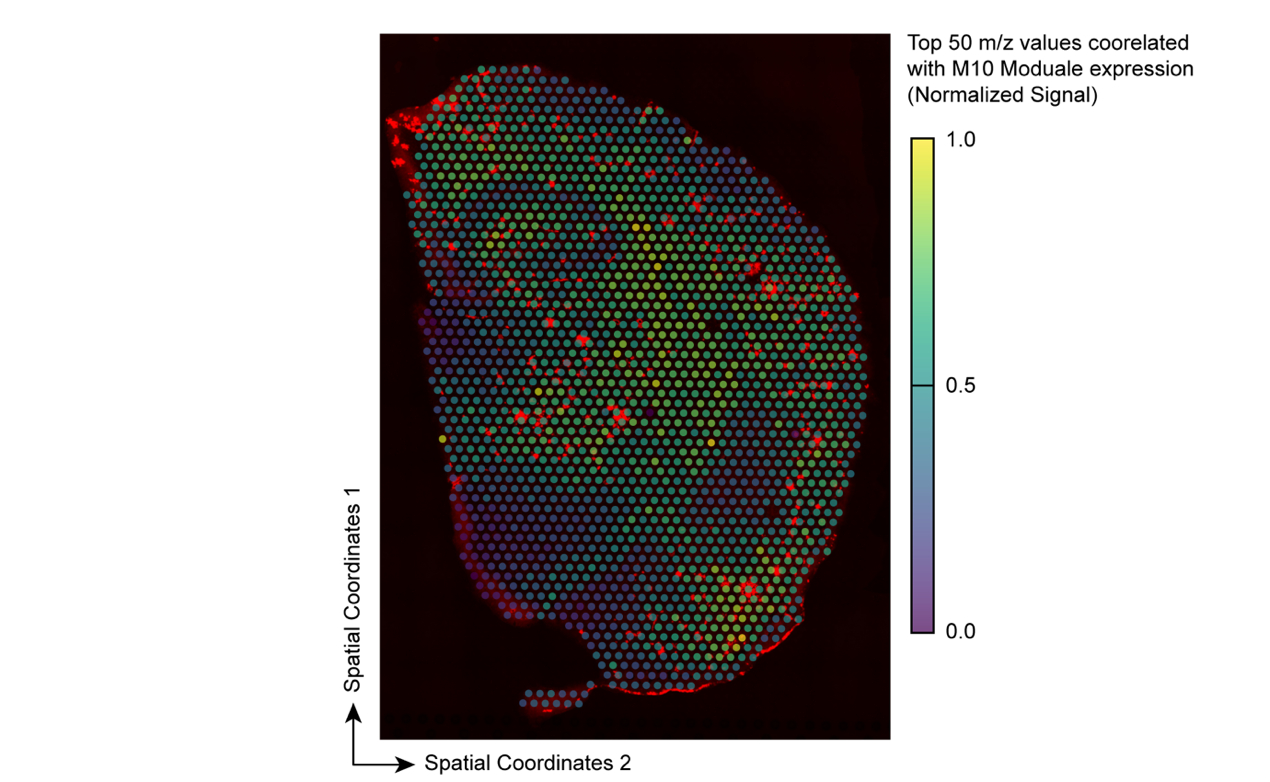

**Supplementary Fig S6. Spatial distribution of the 50 m/z features most correlated with Module 10 genes (WGCNA validation).**

Fluorescent image of stained plaques is overlaid with signal intensity plot based on Visium spots. Maps for all identified modules from WGCNA are provided at DOI: 10.5061/dryad.m37pvmd9h.

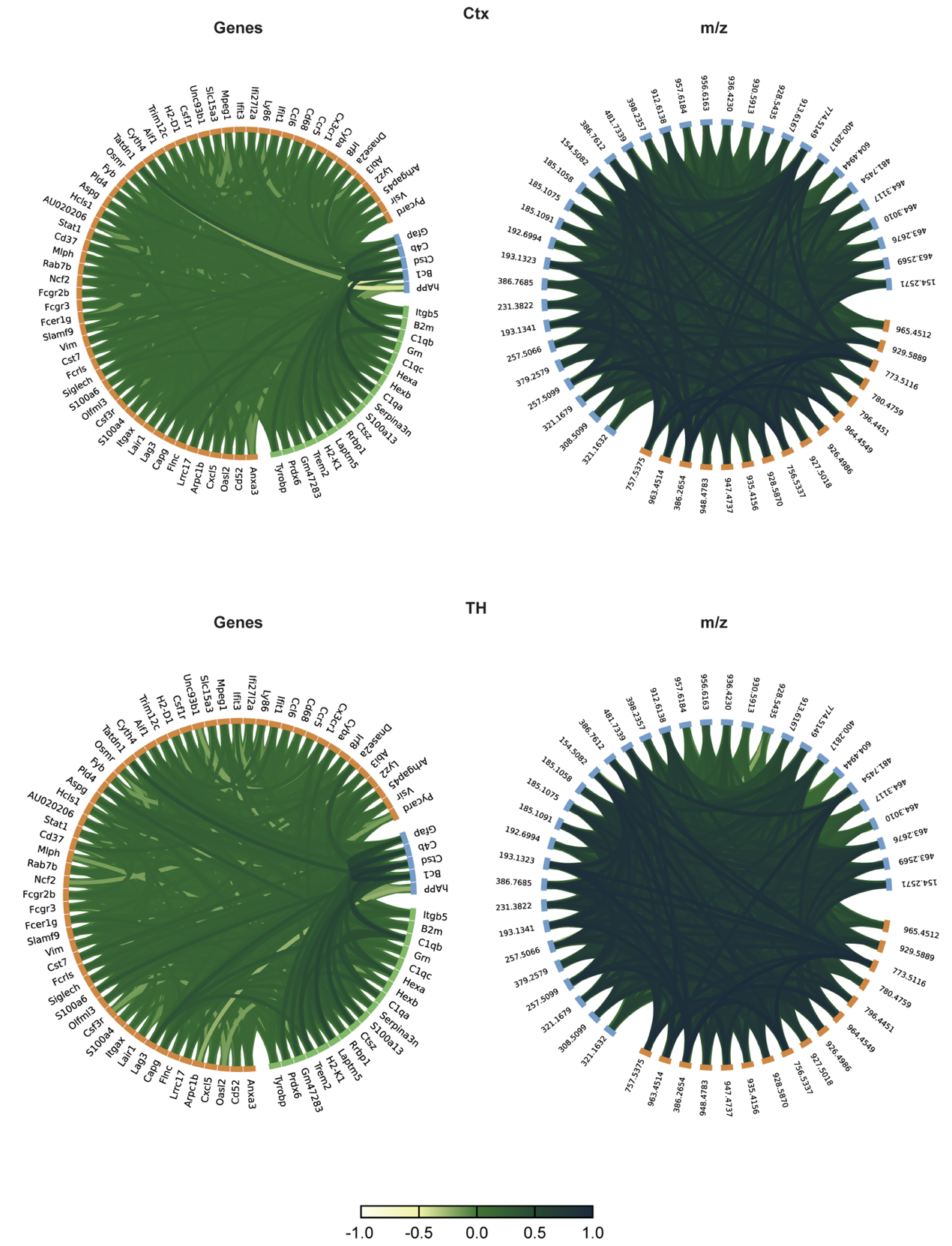

**Supplementary Fig S7. Circos plots of gene or m/z co-correlations for Cortex (Ctx) and Thalamus (TH).** Module 10 genes and the 50 most spatially correlated m/z peaks are displayed in the respective Circos plots. Datasets for all identified brain regions are available at DOI: 10.5061/dryad.m37pvmd9h.

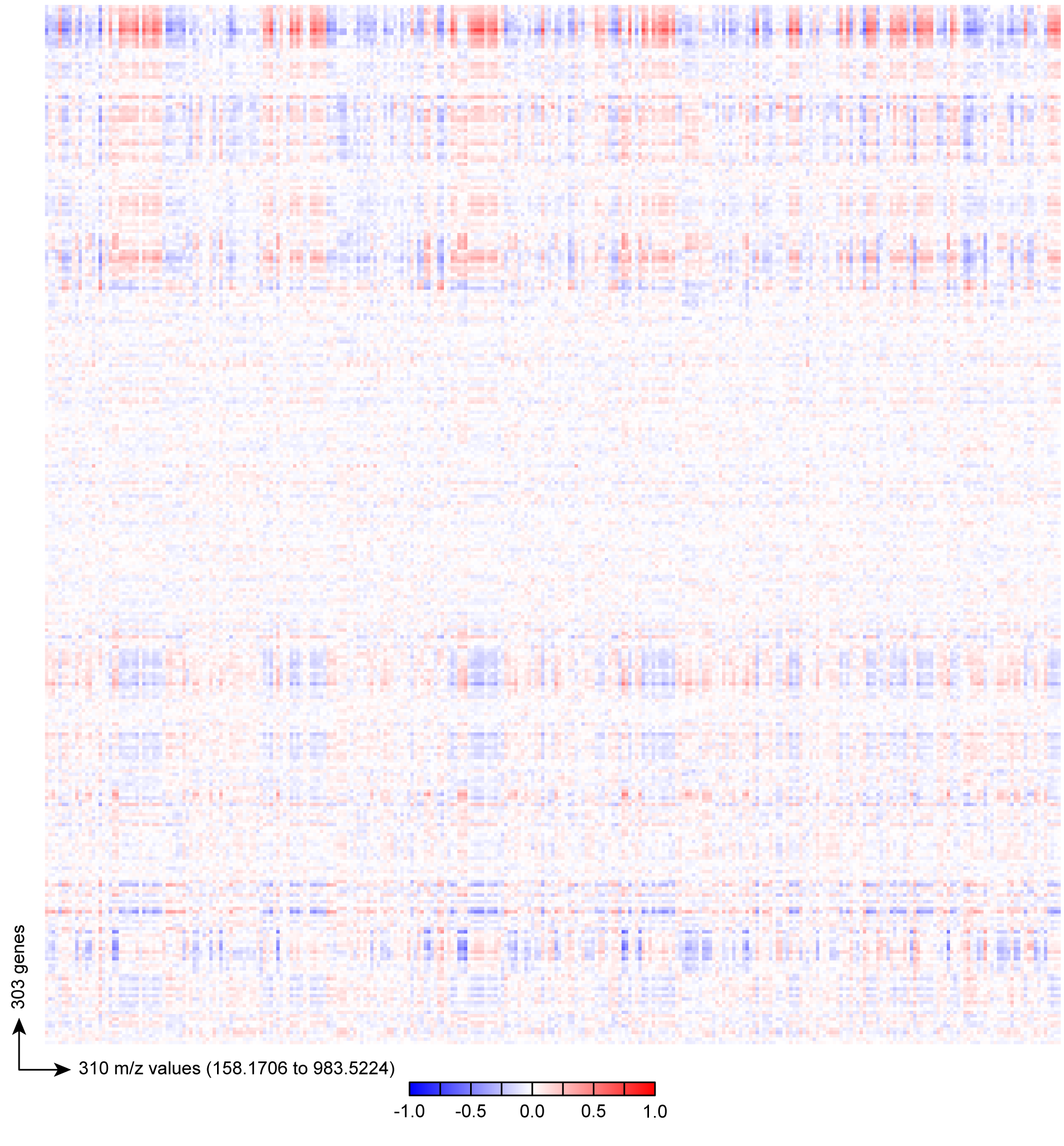

**Supplementary Fig S8. Example of cross‑modal correlation heatmaps between mRNA and mass spectrometry data (Cortex region).** Shown are 303 x 310 feature pairs. Heatmap values refer to correlation coefficient. Datasets for all identified brain regions are available at DOI: 10.5061/dryad.m37pvmd9h.

**Supplementary Data S1**. End-to-end D-iMISTer analysis pipeline with code, parameters, and session info. AVAILABLE AT DOI: 10.5061/dryad.m37pvmd9h.

**Supplementary Data S2**. Table of all differentially expressed genes and metabolites. AVAILABLE AT DOI: 10.5061/dryad.m37pvmd9h.

**Supplementary Data S3**. Table of all significant m/z features with tentative HMDB IDs, adducts, Δppm, fold-change, and ROC AUC values. AVAILABLE AT DOI: 10.5061/dryad.m37pvmd9h.

**Supplementary Data S4**. Violin plots of genes and m/z. AVAILABLE AT DOI: 10.5061/dryad.m37pvmd9h.

**Supplementary Data S5.** Raw RNA and m/z data, with step-by-step instructions for generating Supplementary Fig. S5. AVAILABLE AT DOI: 10.5061/dryad.m37pvmd9h.

**Supplementary Data S6**. KEGG/MetaboAnalyst pathway enrichment reports for each brain region and time point. AVAILABLE AT DOI: 10.5061/dryad.m37pvmd9h.
